# Structure of human sodium leak channel NALCN in complex with FAM155A

**DOI:** 10.1101/2020.07.24.218958

**Authors:** Jiongfang Xie, Meng Ke, Lizhen Xu, Shiyi Lin, Jiabei Zhang, Fan Yang, Jianping Wu, Zhen Yan

## Abstract

NALCN, a sodium leak channel mainly expressed in the central nervous systems, is responsible for the resting Na^+^ permeability that controls neuronal excitability. Dysfunctions of the NALCN channelosome, NALCN with several auxiliary subunits, are associated with a variety of human diseases. Here, we reported the cryo-EM structure of human NALCN in complex with FAM155A, at an overall resolution of 3.1 angstrom. FAM155A forms extensive interactions with the extracellular loops of NALCN that help stabilize NALCN in the membrane. A Na^+^ ion-binding site, reminiscent of a Ca^2+^ binding site in Ca_v_ channels, is identified in the unique EEKE selectivity filter. Despite its ‘leaky’ nature, the intracellular gate is sealed by S6_I_, II-III linker and III-IV linker. Our study establishes the molecular basis of Na^+^ permeation and voltage sensitivity, and provides important clues to the mechanistic understanding of NALCN regulation and NALCN channelosome-related diseases.

## Introduction

Membrane potential across the cell membrane is essential for signal transduction in excitable cells such as neurons and muscle cells (Bean, 2007). In neurons, the resting membrane potential (RMP) is approximately −70 mV, considerably depolarized as compared to the equilibrium potential of K^+^ of −90 mV. The depolarization is mostly attributable to the resting Na^+^ permeability, which is important for the regulation of neuronal excitability (Catterall, 1984; Ren, 2011). The newly identified ion channel NALCN (sodium leak channel, non-selective protein) is primarily expressed in the central nervous system (Lee et al., 1999). It is mainly responsible for the neuronal tetrodotoxin (TTX) -resistant Na^+^ leak conductance and plays a key role in regulating RMP and controlling neuronal excitability (Lu et al., 2007).

NALCN consists of a single polypeptide chain of 24 transmembrane helices (TM) that form four homologous functional repeats connected by intracellular linkers. The topology of NALCN is similar to that of voltage-gated sodium (Nav) channels and voltage-gated calcium (Ca_v_) channels (4×6 TM). Each functional repeat of NALCN contains six TMs (S1-S6), with S1-S4 corresponding to the voltage-sensing domain (VSD) in Na_v_/Ca_v_. S5, S6 and intervening segments, including the pore helices and selectivity filter (SF) from all four repeats, constitute the ion conducting pore. NALCN shares less than 20% identity with any member of Na_v_ and Ca_v_ channels, and it has been classified as a member of a new subclass of the 4×6 TM ion channel family. Compared to Na_v_ and Ca_v_ channels, NALCN contains fewer positively charged residues (arginine or lysine) on S4 segments. Additionally, NALCN has a unique ion selectivity filter (SF) with EEKE residues in the four repeats, which is different from SFs of Na_v_ channels (DEKA) and Ca_v_ channels (EEEE/EEDD) (Lee et al., 1999).

NALCN is associated with several auxiliary subunits, including UNC80, UNC79 and FAM155A. Together they form a large protein complex termed the NALCN channelosome. Notably, these auxiliary subunits are unique to NALCN and share no sequence homology to the auxiliary subunits of any other channel. Both UNC80 and UNC79 are large proteins that contribute to the neuronal localization and stabilization of NALCN in *C. elegans* and *D. melanogaster*, although their own functional domains and subcellular localization are largely unclear (Jospin et al., 2007; Lear et al., 2013; Yeh et al., 2008). UNC80 interacts with NALCN directly and acts as a scaffold protein for UNC79 (Lu et al., 2010; Wang and Ren, 2009). FAM155A, also named NLF-1 in *C. elegans*, was reported to be an endoplasmic reticulum (ER) resident protein that acts as a chaperone to facilitate the folding and promotes axon delivery of NALCN (Xie et al., 2013). In humans, FAM155B, a FAM155A homolog, can functionally substitute for FAM155A, and is likely a component of the NALCN channelosome. In addition, NALCN channelosome activity is regulated by the Src family of tyrosine kinases (SFKs) and several GPCRs such as the M3 muscarinic receptor (M3R) through direct interactions (Swayne et al., 2009). The NALCN channel is also activated by neuropeptides, including substance P and neurotensin, through the SFK-dependent pathway in mouse hippocampal neurons (Lu et al., 2009). How NALCN interacts with and is regulated by its auxiliary subunits remains largely unclear.

NALCN is evolutionally conserved in both invertebrate and vertebrate species. For example, NALCN homolog have been identified in invertebrate species such as snails and *C. elegans*, in which Na_v_ homologues were not found (Lee et al., 1999; Senatore et al., 2013). This indicates that NALCN evolved earlier than Na_v_ channels across species. NALCN is highly conserved in mammals, with the sequence identity of more than 96% among human, mouse, rat, rabbit and bovine. Human NALCN also shares at least 44% identity and 56% similarity with its homologs α1U in *D. melanogaster* and NCA-1/2 in *C. elegans*.

Except for the role in regulating neuronal excitability, NALCN is also important in many fundamental physiological processes, such as motor function, pain sensitivity and circadian rhythm. For example, NALCN mutant mice die within 24 hours after birth due to disrupted respiratory rhythm (Lu et al., 2007). Overexpression of NALCN in highveld mole-rat leads to ablating detection of painful substances by nociceptors (Eigenbrod et al., 2019). Reduced NALCN expression in *Drosophila* leads to changes in behavioral circadian rhythms and sensitivity to anesthetics (Lear et al., 2005; Nash et al., 2002). In humans, NALCN variants are linked to a variety of diseases, including congenital contractures of limbs and face, hypotonia and developmental delay (CLIFAHDD)(Bend et al., 2016; Chong et al., 2015; Fukai et al., 2016), psychomotor retardation and characteristic facies (IHPRF) (Al-Sayed et al., 2013; Angius et al., 2018; Bramswig et al., 2018), infantile neuroaxonal dystrophy (INAD) (Koroglu et al., 2013), cervical dystonia, schizophrenia and bipolar disorder (Askland et al., 2009; Mok et al., 2014; Wang et al., 2010). There are also many diseases related to the dysfunction of the auxiliary subunits of NALCN (Shamseldin et al., 2016; Stray-Pedersen et al., 2016).

Despite its physiological and pathological importance, the biophysical properties of the voltage sensitivity and ion selectivity of NALCN are still under debate. Electrophysiological studies on NALCN have been performed in heterologous expression systems such as HEK293 cells, *X. laevis* oocytes and the neuronal cell line NG108-15 (Bouasse et al., 2019; Chua et al., 2020; Lu et al., 2007). NALCN expressed in HEK293 cells was reported to exhibit a linear current-voltage relationship, suggesting that NALCN is a voltage-independent ion channel. The same study also reported that NALCN is nonselective and permeable to Na^+^, K^+^ and Ca^2+^ (Lu et al., 2007). Other studies suggested that the VSDs of NALCN can exhibit a broader range of gating behaviors. Co-expression of NALCN, UNC79, UNC80, and FAM155A resulted in voltage dependent NALCN current (Bouasse et al., 2019; Chua et al., 2020). Finally, NALCN was also reported to be only selective for monovalent cations and to be blocked by extracellular divalent cations (Chua et al., 2020).

To better understand the functional properties and mechanisms of NALCN, we sought to determine the structure of NALCN in complex with its auxiliary subunits. Here, we report the cryo-EM structure of human NALCN in complex with FAM155A at an overall resolution of 3.1 Å. Our structure along with electrophysiology and molecular dynamic (MD) simulation data, provide unprecedented insights in understanding the structure, function and regulation of the NALCN channelosome.

## Results

### Functional characterization of NALCN by electrophysiology

We employed patch-clamp recordings in HEK293 cells to characterize the electrophysiological properties of NALCN. The current elicited in response to voltage steps was very small and appeared to be indistinguishable from the mock when NALCN channel was expressed alone in HEK293 (Figure S1A). By contrast, co-expression of NALCN with UNC79, UNC80 and FAM155A dramatically enhanced the current, which was largely inhibited by 1 mM verapamil, implying that the current measured was mainly mediated by NALCN (Figure S1B). We also clearly recorded the kinetics of voltage sensitive NALCN current: hyperpolarization voltages elicited large inward current with inactivation, while depolarization voltages activated current with an exponential course before reaching in the steady state (Figure S1B). These observations indicate that the auxiliary subunits are essential for a functional NALCN (Figure S1A), consistent with previous studies (Bouasse et al., 2019; Chua et al., 2020). The electrophysiological properties of NALCN suggested that it tends to maintain the RMP in a manner reminiscent the Le Chatelier’s principle in chemical equilibria. In comparison, Na_v_ channels always show rapid current activation and inactivation for fast initiating action potential (Catterall, 2000), while certain Kv such as KCNQ channels show slow activation kinetics to avoid fast repolarization (Folander et al., 1990).

### Structure determination of human NALCN-FAM155A complex

Based on our functional characterizations, we focused on the co-expression of the four components in HEK293 cells. Unfortunately, we were unable to get a stable complex of the four components. Instead, a stable and homogeneous subcomplex of NALCN and FAM155A was obtained (Figure S2A). The peak fractions from gel filtration purification, were pooled and concentrated to about 8.5 mg/ml for cryo-EM sample preparation. The EM images were collected on a Titan Krios electron microscope operating at 300 kV and equipped with a K3 direct detector and a Gif Quantum energy filter. After a few rounds of two-dimensional (2D) and 3D classifications, about sixty-five thousand particles were selected, which yield a final reconstruction map with an overall resolution of 3.1 Å (Figure 1, Table S1, Figure S2, S3). Local masks of the extracellular and intracellular regions were applied to further improve the local resolutions to 2.9 Å and 3.1 Å, respectively.

**Figure 1.**
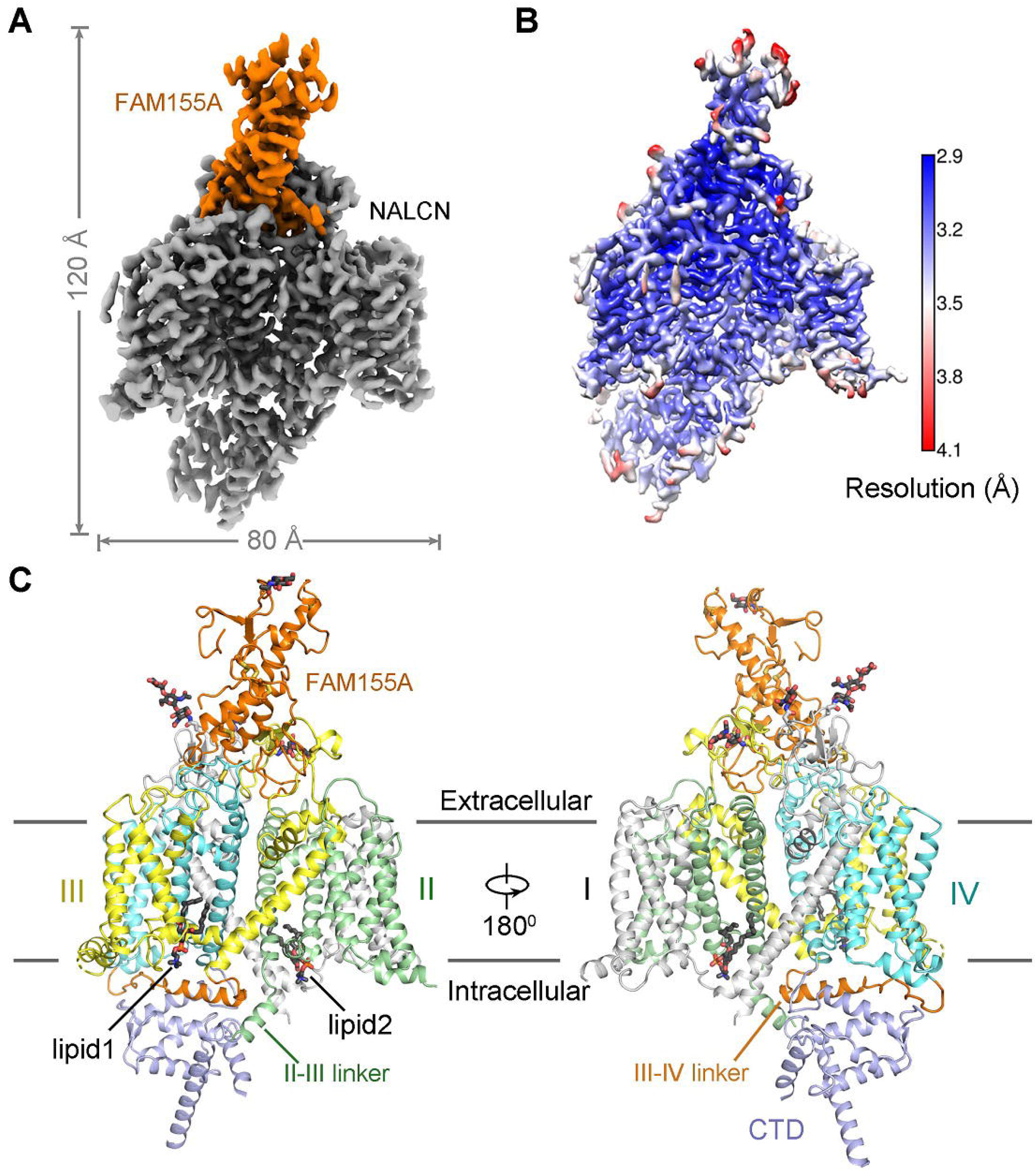
Cryo-EM structures of the human NALCN in complex with FAM155A. **(A)** Cryo-EM map of the human NALCN in complex with FAM155A. NALCN and FAM155A are colored in gray and orange, respectively. The map was generated in ChimeraX (Goddard et al., 2018). **(B)** Local resolution map estimated by cryoSPARC (Punjani et al., 2017) and generated in Chimera (Pettersen et al., 2004). **(C)** Overall structure of NALCN-FAM155A complex presented in two side views. The protein structure was shown in cartoon. NALCN was domain colored with repeat I, II, III, IV and CTD in light gray, green, yellow, cyan and slate, respectively. The III-IV linker and FAM155A are colored in orange. Two lipid molecules bound in the inner leaflet of membrane and the sugar moieties at the glycosylation sites are shown in stick. The color schemes are applied in all figures. All structure figures were prepared with PyMol (DeLano, 2002).

The density of the final reconstruction is of high quality with evenly distributed estimated local resolution. Most side chains were clearly resolved that allow for accurate model building (Figure 1A, 1B, Figure S4, S5). The model of NALCN was built with a homology model based on Ca_v_1.1 (PDB: 5GJV), whereas FAM155A was built *de novo*. In total, 1295 and 182 residues were built for NALCN and FAM155A, respectively. The complex structure was confirmed by cross-linking mass spectrometry analysis (Figure S6). Three glycosylation sites (Asn210, Asn216 and Asn1064) and one glycosylation site (Asn217) were identified on NALCN and FAM155A, respectively (Figure 1C, Figure S5C). Four disulfide bonds in the extracellular loops (ECL) of NALCN and six in FAM155A were recognized (Figure 1C, Figure S5D). The glycosylation sites and disulfide bonds in turn verified the accuracy of the structure.

The overall structure of NALCN resembles those of eukaryotic Na_v_ and Ca_v_ channels (Figure 2). We select human Na_v_1.4 (PDB: 6AGF) and rabbit Ca_v_1.1 (PDB: 6JP5) as representatives of Na_v_ and Ca_v_ channels for comparisons with NALCN (Pan et al., 2018; Zhao et al., 2019). The structures of human Na_v_1.7 (PDB: 6J8I), rat Na_v_1.5 (6UZ3) and Na_v_PaS (6A95) from American cockroach were also picked for specific discussions (Jiang et al., 2020; Shen et al., 2018; Shen et al., 2019). The structures of NALCN can be superimposed to the α subunit of Na_v_1.4 and α1 subunit Ca_v_1.1 with root-mean-square deviation (r.m.s.d) of 2.99 Å over 760 Cα atoms and 3.894 Å over 905 Cα atoms, respectively. Except for the four homologous repeats, a C-terminal domain (CTD) after repeat IV, which was observed in the structures of Ca_v_1.1 and Na_v_PaS but not in human Na_v_ channels, was also clearly resolved in the structure of NALCN (Figure 1C). NALCN has no N-terminal domain (NTD) before repeat I, which was always observed in human Na_v_ channels.

**Figure 2.**
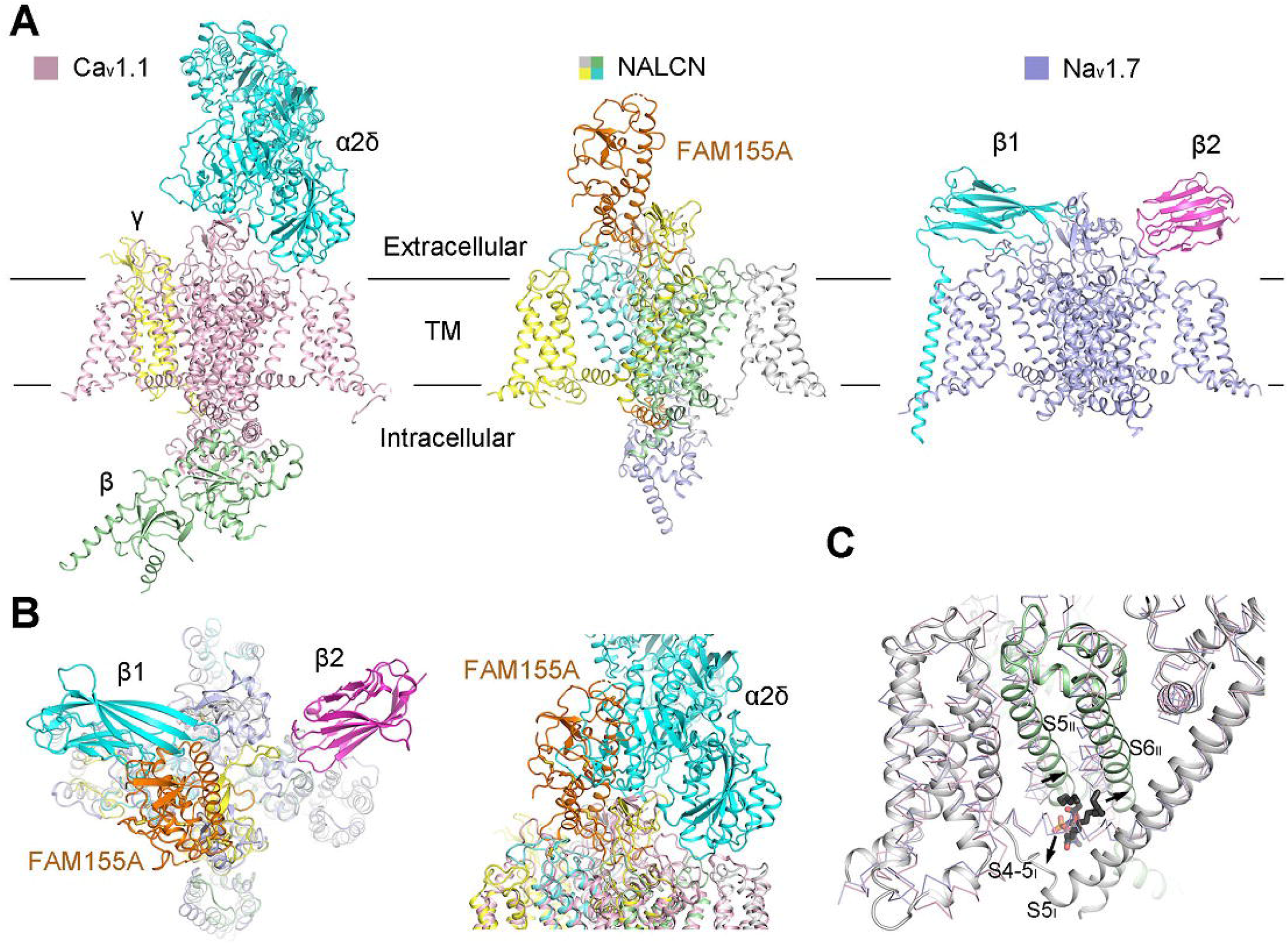
Structure comparisons between NALCN and Na_v_/Ca_v_ channels. **(A)** Overall structures of human NALCN, Na_v_1.7 (PDB: 6J8I) and rabbit Ca_v_1.1 complex (PDB: 6JP5). The domain color scheme of NALCN is the same as in Figure 1. The ion conducting subunit of Na_v_1.7 and Ca_v_1.1 are colored slate and light pink, respectively. The auxiliary subunits β1 and β2 of Na_v_1.7 are colored cyan and magenta and the auxiliary subunits α2δ, β, γ of Ca_v_1.1 are colored cyan, yellow and green, respectively. **(B)** Superimposition of the structures between NALCN-FAM155A and Na_v_1.7 complex or Ca_v_1.1 complex. FAM155A adopt a unique binding site on NALCN that differs from the auxiliary subunits of Na_v_/Ca_v_ channels. The structure alignments is based on the ion conducting subunit of each complex. Na_v_1.7 is shown in an extracellular view, and Ca_v_1.1 is shown in a side view. **(C)** Structure comparisons of the Repeat I and repeat II among NALCN, Na_v_1.7 and Ca_v_1.1. NALCN is shown in cartoon and Na_v_1.7 and Ca_v_1.1 are shown in ribbon. The S5_I_ in NALCN is extended into cytosol. The linker between S4 and S5 segments (S4-5 linker) in repeat I of NALCN is a loop instead of a helix in Na_v_1.7/Cav1.1. An identified lipid molecule in NALCN occupies in a position that correspond to the S4-5 helix in Na_v_1.7/Cav1.1. Local structural deviations around the lipid between NALCN and Na_v_1.7/Cav1.1 are indicated by black arrows.

NALCN has several unique features that are distinct from both Na_v_ and Ca_v_ channels. In particular, two lipids, lipid1 near repeat I and lipid2 near repeat III, were found to bind NALCN at the inner membrane side (Figure 1C). Lipid2 resides in a cavity surrounded by S3, S4 and S4-5 of repeat III, while lipid1 occupies the cavity surrounded by S4_I_, S5_I_, S6_I_, S5_II_, S6_II_ and S4-5_I_ (Figure 2C, Figure S4B). Notably, lipid1 sits in a position that occupied by S4-5 linker in Na_v_/Ca_v_ structures (Figure 2C). This may have caused the conformational change of the S4-5_I_ linker from a helix to a loop in NALCN. In addition, a short helix designated as the II-III linker, that was connected to S6_II_ by a short loop was firstly observed in the structure of NALCN (Figure 1C).

### Interactions between NALCN and FAM155A

Human FAM155A contains 456 amino acids and residue 192-382 were resolved in our structure except for a short loop from residue 250 to 258 (Figure 3A). The N-terminal and the C-terminal regions of FAM155A, including two predicted TM helices, were not resolved in current structure, probably because of their intrinsic flexibility. The resolved structure of FAM155A contains two lobes, designated as N-lobe and C-lobe, stabilized by six disulfide bonds. The N-lobe consists of three short β strands and two short α helices (H1-H2), while the C-lobe contains four α helices (H3-H6) (Figure 3B). The overall structure of FAM155A is different from any auxiliary subunit in Na_v_ or Ca_v_ channels (Figure 2B, Figure S7A). A search of the protein data bank using the DALI server (Holm, 2019) revealed that the C-lobe of FAM155A contains a fold (similarity Z-score 4.3) similar to the structure of the frizzled-like cysteine-rich domain (CRD) of receptor tyrosine kinase MuSK (PDB: 3HKL), which had 84 Cα atoms aligned to FAM155A with a r.m.s.d. of 2.8 Å (Figure S7B). Superimposition of the CRD domain of MuSK on FAM155A shows that H3, H5 and H6 align quite well. Moreover, three disulfide bonds are structurally conserved between the two structures (Figure S7B).

**Figure 3.**
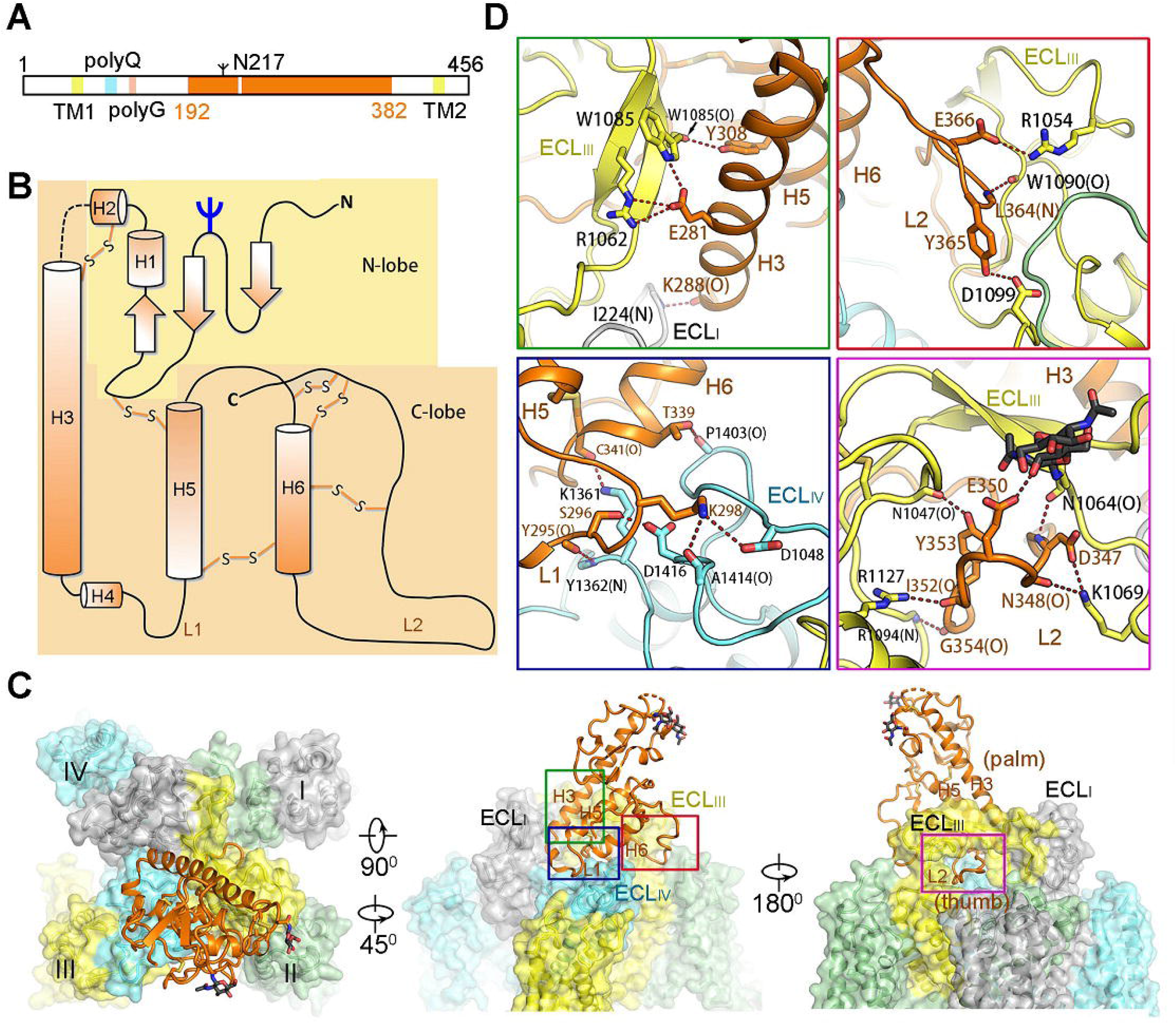
Specific interactions between NALCN and FAM155A. **(A)** 1-dimensional (1D) schematic view of FAM155A. The region that was resolved in current structure was colored in orange. The position of glycosylation site, two predicted transmembrane helices, polyQ and polyG motifs were labeled. **(B)** 2D topology structure of FAM155A. The glycosylation site and six disulfide bonds were labeled. Six helices were named as H1-H6, respectively. The loop connecting H4 and H5 and the loop after H6 were designated L1 and L2, respectively. **(C)** Interaction interface between NALCN and FAM155A. NALCN is shown in surface and FAM155A is shown in cartoon. Details in the rectangle boxes are presented in panel D. **(D)** Zoom in views of each interaction interface as shown in panel C. The electrostatic interactions are indicated by red dash lines. The residues from NALCN and FAM155A are labeled black and orange, respectively. Please also refer to Table S2 for a summary of interaction between NALCN and FAM155A.

Only the C-lobe of FAM155A is involved in the interaction with NALCN. FAM155A sits above the pore domain of repeat IV of NALCN and form extensive interactions with the ECLI, ECLIII and ECLIV of NALCN (Figure 1C, 3C, Table S2). The loop in FAM155A connecting H4 and H5 and a hairpin loop after H6 that is involved in binding to NALCN are designated L1 and L2, respectively (Figure 3B). Notably, L2 of FAM155A inserts deeply into a dome formed by ECLIII of NALCN. The H3, H5, H6 helices and L2 of FAM155A resemble the palm and thumb of a hand, respectively, holding ECLIII of NALCN tightly (Figure 3C). Detailed analysis of the interaction interfaces revealed extensive salt bridges and hydrogen bonds between NALCN and FAM155A (Figure 3D, Table S2). Unexpectedly, the sugar moiety linked to Asn1064 on ECLIII, also contributes to the interaction, by forming a hydrogen bond with E350 on FAM155A (Figure 3D). Electrostatic surface analysis revealed that two surfaces between NALCN and FAM155A are electrically complementary, ensuring a favorable interaction stability (Figure S7C). Sequence alignment among NALCN and Na_v_/Ca_v_ channels revealed that the interface residues in NALCN, including Asn1064, are not conserved among other Na_v_/Ca_v_ channels, explaining the high binding specificity of FAM155A towards NALCN, but not other channels (Figure S8). The interface between FAM155A and NALCN is very different from those between the ion conducting α subunit of Na_v_ and Ca_v_ channels and their auxiliary subunits (Figure 2B). The key residues in the interaction interface are highly conserved among NALCN and FAM155A proteins from different species, indicating that the interaction mode between NALCN and FAM155A is evolutionarily conserved. Furthermore, the key residues mediating complex formation in FAM155A are also conserved in FAM155B, another subtype protein of the FAM155 family (Figure S9). Therefore, FAM155B may also be able to form a stable complex with NALCN *in vivo*. This notion is supported by the finding that FAM155A can be functionally substituted by human FAM155B (Chua et al., 2020).

As our structure was captured as a subcomplex between NALCN and FAM155A, we then asked whether co-expression of NALCN and FAM155A only was also able to recapitulate measurable NALCN current. Interestingly, we have readily recorded large current in response to voltage steps when NALCN and FAM155A was co-transfected (Figure S1C). The current-voltage curve of NALCN co-expressed with either all three auxiliary subunits or with FAM155A only was non-linear (Figure S1D). Compared with NALCN co-expressed with three auxiliary subunits, NALCN with FAM155A exhibited similar voltage sensitivity in current as the apparent gating charge (Z_app_) was similar (Figure S1D, Z_app_: 1.25 ± 0.55 e_0_ and 1.22 ± 0.18 e_0_, respectively). However, the G-V curve of NALCN co-expressed with three auxiliary subunits was largely shifted towards the hyperpolarization voltage as compared to NALCN with FAM155A (V_1/2_: −44.66 ± 1.91 mV and −65.07 ± 2.11 mV, respectively) (Figure S1E). Moreover, we observed no inactivation of the steady state current by setting the prepulse potential to different levels when NALCN channel was co-expressed with auxiliary subunits (Figure S1F and S1G), which is consistent with the physiological role of NALCN being a leaky sodium channel.

### Structural basis for the voltage dependence of NALCN

Our structural study has provided an opportunity to understand the voltage sensitivity of NALCN. Detailed analysis of the voltage sensing domains (VSDs) of NALCN suggest that they possess several key features shared among functional VSDs in Na_v_/Ca_v_ channels. For instance, each VSD preserves the gating charges (GCs) in a pattern of occurring every three amino acids in the S4 segments. As usual, we define the position of last gating charge in each S4 as R6 (Wu et al., 2016). NALCN has two to four GCs in each repeat distributed at positions R2-R6 (Figure 4A). The last GC of repeat IV is at a position that has one residue shift to R6, a phenomenon also observed in other channels (Wu et al., 2016). Moreover, the residues of charge transfer center (CTC) consists of a negatively charged residue (An2) on S2, an aromatic occluding residue (F) on S2, and a negatively charged residue on S3 are highly conserved in NALCN except for the occluding residue on VSDIII. The An1 sites that are seven residues ahead of the occluding residues on S2 segments of NALCN are mainly negatively charged or polar residues, similar to those of Na_v_/Ca_v_ channels (Figure 4A, B). In addition, previous structural studies on Na_v_ channels identified several negatively charged or polar residues on S1 that may also play roles in gating charge transfer (Pan et al., 2018). These residues were also observed on NALCN (Figure 4A). When the four VSDs are superimposed relative to An1 and CTC, all R4 residues are above the occluding residues on S2, reminiscent of an up or depolarized state of the four VSDs (Figure 4B, Figure S10).

**Figure 4.**
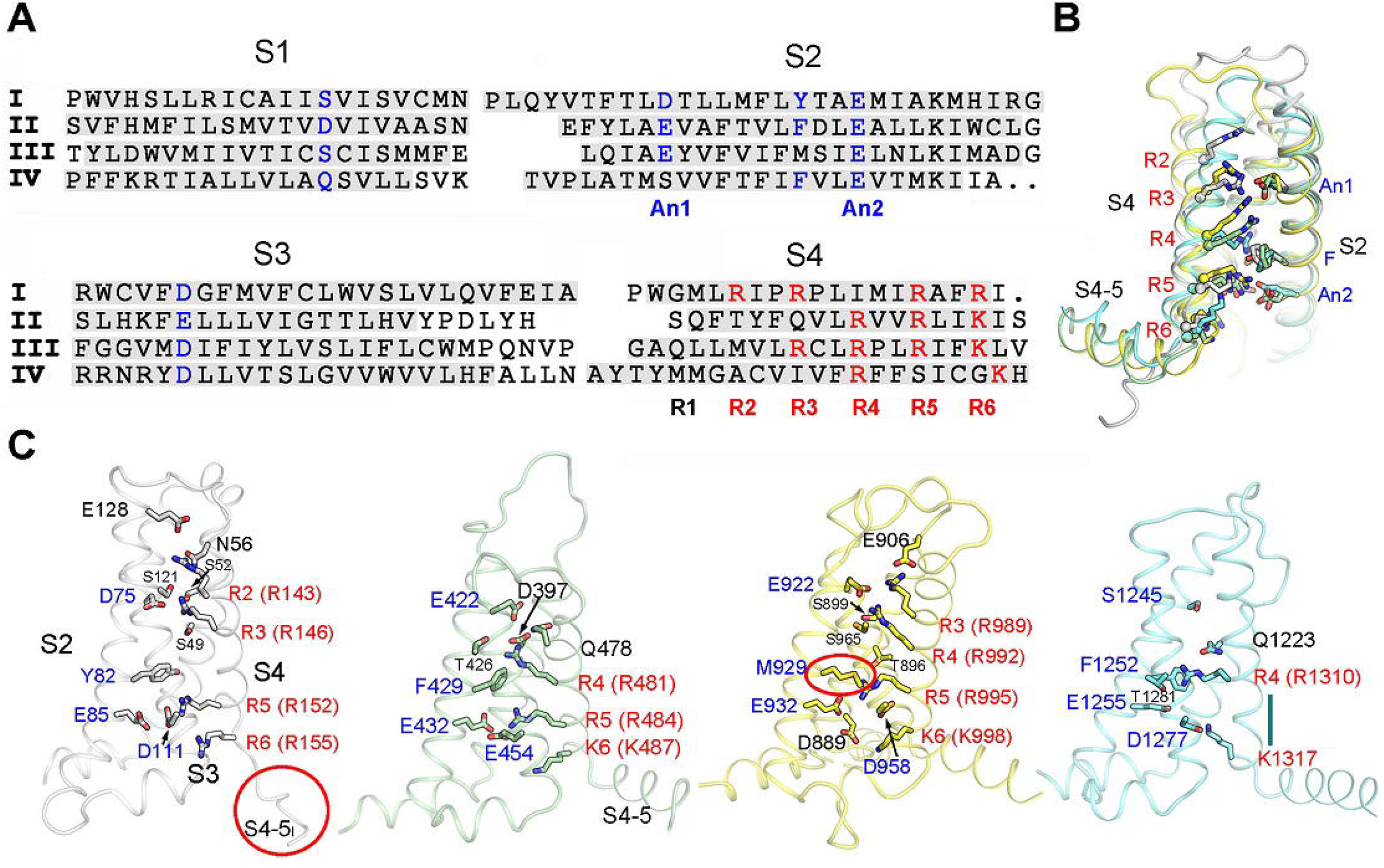
Structural features of the VSDs of NALCN. **(A)** Structural-based sequence alignment of the four VSDs. The boundaries for the S1 to S4 segments are shaded in gray. The gating charges (GCs) in the S4 helices are colored in red. The residues that correspond to the CTC on S2 and S3 helices as well as the polar or acidic residues on S1 that coordinates GCs are highlighted in blue. **(B)** Structure comparison of the four VSDs. The four VSDs are superimposed relative to CTC and An1 on S2. For visual clarity, the S1 segments were omitted. All four VSDs adopt up or depolarized state. **(C)** Structural details of each VSD. The GCs, CTC, and the polar or acidic residues participating in GCs coordination are shown in sticks. The GCs are labeled in red. The S4-5_I_ loop and non-conserved aromatic residues in the CTC of repeat III are highlighted by red circles. The short 3_10_ helix turn in repeat IV is indicated by a straight line.

The voltage sensitivity of NALCN, however, seems to be weaker than that of Na_v_/Ca_v_ channels (Figure S1). Several unique structural features of NALCN may lead to its relative weak voltage sensitivity. First, the occluding residue on S2 in repeat III is replaced by a methionine (Met929), instead of a conserved phenylalanine or tyrosine (Figure 4C). The occluding residue of the CTC is crucial for the gating transfer during voltage sensing. A previous study shows that mutations of the occluding residue from bulky hydrophobic residue to methionine on a voltage-gated potassium channel diminish its voltage sensitivity (Tao et al., 2010). Second, the S4 of VSDs are usually formed as 3_10_ helices, whereas in repeat IV the S4 segment largely relax to regular α helix. The only 3_10_ helix turn in S4_IV_, which is at position of R5, is a serine instead of a GC. These structural observations are also consistent with reported experimental data. For example, it was reported that repeat III and IV contribute little to the voltage sensitivity of NALCN, probably due to their lack of an occluding residue on S2, and lack of GCs and 3_10_ helices on S4, respectively. In contrast, neutralization of the GCs R146+R152 in repeat I or R481+R484 or R481+K487 in repeat II lead to significantly changes in voltage sensitivity (Chua et al., 2020). Third, the upper gating charges (R2-R4) in repeat I-III form more extensive hydrogen bonding interactions with surrounding negatively charged or polar residues, compared with their counterparts in Na_v_/Ca_v_ channels (Figure 4C). During the gating charge transfer process, the GCs need to break old interactions with surrounding residues and form new interactions with other residues, accompanying with a sliding movement along the S4 segment. The extensive interactions of the GCs and surrounding residues in NALCN may need the VSDs to overcome a higher energy barrier to undergo conformational changes thus resulting in a weaker voltage sensitivity. Fourth, the S4-5_I_ linker unexpectedly forms a flexible loop instead of a juxtamembrane helix seen in other repeats and other reported Na_v_/Ca_v_ structures. Meanwhile, S5_I_ is extended and elongated to the cytoplasmic region with three helix turns. The unique structure of S4-5_I_ and S5_I_ results in a cavity that accommodate a lipid, that may contribute to the regulation of NALCN (Figure 2C, Figure S4B). As S4-S5 is supposed to be a key transducer from VSDs to the ion conducing pore during voltage sensing gating in VGICs (Yan et al., 2017), its relaxation to a flexible linker may hinder the effective transduction of electromechanical coupling of repeat I in NALCN. Altogether, these structural variations in NALCN may contribute to its weak voltage sensitivity.

### EEKE selectivity filter in NALCN

Like all reported Na_v_/Ca_v_ structures, the pore domain of NALCN is formed by S5 and S6 helices of the four repeats and the selectivity filter (SF) of NALCN was supported by two pore helices (P1 and P2) that intervene between S5 and S6 (Figure 5A). The key residues in the SF of NALCN responsible for the ion selectivity are E280/E554/K1115/E1389 (EEKE), distinct from DEKA in Na_v_ channels and EEEE/EEDD in Ca_v_ channels. Sequence alignment of the P1-SF-P2 among NALCN, Ca_v_1.1 and Na_v_1.4 reveals an invariant tryptophan in the first residue of P2 and a highly conserved residue (Thr/Ser/Cys) in the last residue of P1, while other residues are not very conserved (Figure 5B). The overall structure of P1-SF-P2 in NALCN is more closely related to that of Ca_v_1.1 than to that of Na_v_1.4, with r.m.s.d. of 0.848 Å over 110 Cα atoms and 1.602 Å over 107 Cα atoms, respectively. Both the pore helices and the SF of NALCN align well with those of Ca_v_1.1, whereas the P2s are not aligned well in superimposition between NALCN and Na_v_1.4 (Figure 5C). Despite its structural similarity with Ca_v_ channels, NALCN was reported to be mainly selective for monovalent cations and could be blocked by extracellular divalent cations (Chua et al., 2020). In addition, NALCN is responsible for the background sodium leak conductance in hippocampal neurons, indicating that NALCN is selective for Na^+^ ion *in vivo* (Lu et al., 2007).

**Figure 5.**
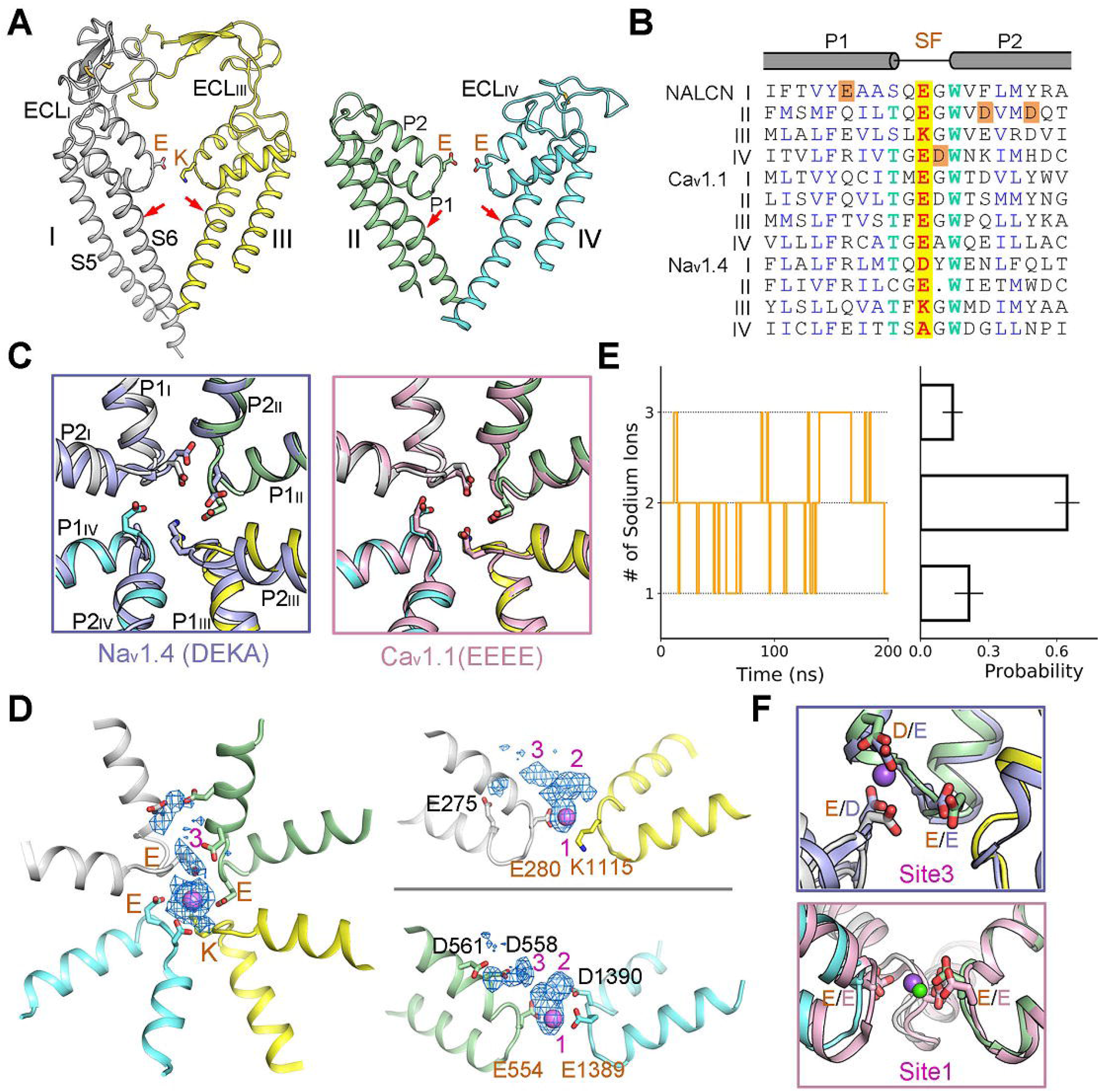
The EEKE selectivity filter of NALCN. **(A)** Overall structure of the pore domain of NALCN. Side views of the pore domain from the diagonal repeats are shown. The EEKE residues in the selectivity filter (SF) are shown in stick. Repeat I and repeat III adopt large extracellular loops, reminiscent that of Ca_v_1.1. A π helix turn was observed in each S6 segment and was indicated by red arrows. **(B)** Sequence alignment of the SF and the connecting pore helices among human NALCN, Ca_v_1.1 and Na_v_1.4. The critical EEKE (E280, E554, K1115, and E1389) residues in NALCN and the corresponding residues in Ca_v_1.1 and Na_v_1.4 are shaded yellow. The invariant Trp residues and the highly conserved Thr residues are colored light green. Abbreviations: A, Ala; C, Cys; D, Asp; E, Glu; F, Phe; G, Gly; H, His; I, Ile; K, Lys; L, Leu; M, Met; N, Asn; P, Pro; Q, Gln; R, Arg; S, Ser; T, Thr; V, Val; W, Trp; Y, Tyr. **(C)** Structure comparisons of the SF between NALCN and Na_v_1.4 (*left*) or Ca_v_1.1 (*right*). The critical residues in the SF are shown in stick. **(D)** Top view and two side views of sodium probability density generated from pore domain equilibration. Three potential Na^+^ binding site were identified and labeled in purple. The most stable binding site (Site1) is signposted by a Na^+^ ion (magenta sphere). Please also refer to Movie S1 for the illustrative trajectory of equilibration. **(E)** Ion numbers within 4 Å of SF residues (EEKE) through time (*left*) and the probability statistics (*right*). The statistics represent the results of three independent simulations. **(F)** Structural comparison of Site 3 (EED motif) in NALCN and the Na^+^ binding site (DEE motif) in Na_v_1.4 (*top*) and of Site1 in NALCN and a Ca^2+^ binding site in Cav1.1 (bottom). Na^+^ and Ca^2+^ are shown as spheres in purple and green, respectively.

To better understand Na^+^ binding and selectivity in the SF, we performed canonical MD simulation to investigate the interaction of membrane-embedded pore domain in 150 mM NaCl. In three independent trials of 200 ns equilibration, we observe three potential ion binding sites, designated Site1-3, in proximity to SF (Figure 5D). In addition to the glutamic acid residues in SF, the Na^+^ binding sites are formed by several negatively changed residues from the pore helices, including E275, D558, D561 and D1390. Among the three binding sites, Site1 is most stably located at the middle of SF, while the other two are transient and presumably cation attractants (Figure 5D, Movie S1). The simulation results were reproducible in three parallel runs. Probability statistics suggest that there is an average of two Na^+^ ions within 4 Å range of the SF residues through time (Figure 5E). Notably, Site3 formed by E280/E554/D558 (designated EED site) is spatially close to the Na^+^ binding site formed by a DEE motif in Na_v_ channels (Figure 5F) (Pan et al., 2018). However, the sidechains of the EED motif in NALCN are not as closely spaced as the DEE site in Na_v_ channels and do not form a favorable Na^+^ binding site. On the other hand, Site1, the most occupied binding site, is spatially close to a putative Ca^2+^ binding site in Ca_v_1.1 and Ca_v_3.1, suggesting that NALCN and Ca_v_ channels share similar ion selectivity mechanisms (Wu et al., 2016; Zhao et al., 2019) (Figure 5F). The replacement of glutamic acid (E) or aspartic acid (D) with lysine (K) in repeat III from Ca_v_ channels to NALCN has shifted the ion selectivity from Ca^2+^ to monovalent ions. Like Na_v_ channels, the lysine in NALCN favors Na^+^ ion and blocks Ca^2+^ permeation through the selectivity filter. Meanwhile, the EEKE SF may still preserve the ability to bind Ca^2+^ as the structure of SF in repeat I/II/IV is almost identical to that of Ca_v_ channels (Figure 5C). In this sense, extracellular Ca^2+^ may compete with Na^+^ in binding to the SF, explaining why Ca^2+^ is a blocker of NALCN as shown by previous study (Chua et al., 2020).

### Closed intracellular gate of NALCN

The ion permeation path below the SF in NALCN is enclosed by the S6 tetrahelical bundle, a structural feature that is shared in all reported Na_v_/Ca_v_ structures (Figure 5A, 6A). However, due to the existence of FAM155A, ions can only enter the permeation path from one side, that is above the pore domain of repeat II (Figure 6A, *left*). We used HOLE to calculate the radius along the permeation path and compared it with Na_v_1.4 and Ca_v_1.1, whose intracellular gates are in open and closed state, respectively. Surprisingly, although NALCN is supposed to conduct a ‘leak’ current, the intracellular gate is closed, even tighter than the closed Ca_v_1.1. The narrowest region, sealed by two layers of hydrophobic residues Val, Ile and Leu, is about 10 Å in length along the permeation path with a radius less than 1 Å (Figure 6A, *right*). Notably, the lower gate is fully blocked by the III-IV linker, causing the permeation path to enter the cytosol only from the side. Detailed analysis reveals that the III-IV linker is stabilized by extensive hydrogen bonds or polar interactions with S6_I_ and the II-III linker (Figure 6B). Most of the involved residues are not conserved among Na_v_/Ca_v_ channels, indicating that the local interactions among S6_I_, the II-III liner and the III-IV linker are highly specific to NALCN, which implies distinct gating mechanism of NALCN (Figure S7).

**Figure 6.**
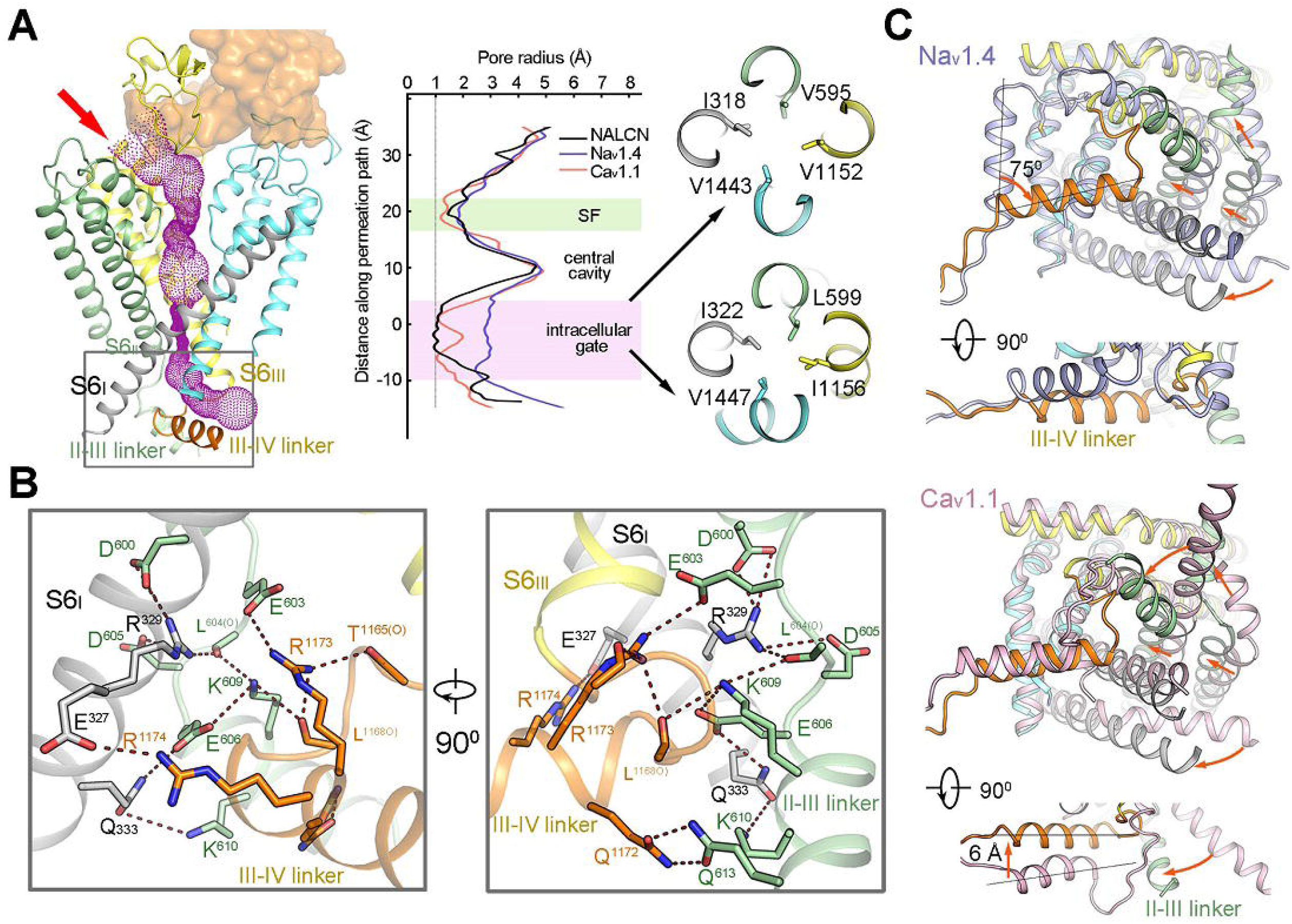
A closed intracellular pore. **(A)** Ion permeation path of NALCN. The pore region of NALCN is shown in cartoon and FAM155A is shown in surface. The ion permeation path calculated by HOLE (Smart et al., 1996) is illustrated by purple dots on the left. The extracellular ion entrance is indicated by red arrow. The details in the rectangle region is shown in panel B. The corresponding pore radii along the permeation path of human NALCN (black), Na_v_1.4 (slate), and Ca_v_1.1(pink) are compared in the middle. Two narrowest layers of the intracellular gate are presented in extracellular views on the right. **(B)** Zoom in views of the intracellular gate. The electrostatic interactions are indicated by red dash lines. The residues from S6_I_, II-III linker, and III-IV linker are colored black, green, and orange, respectively. **(C)** Pore structure comparisons between NALCN and Na_v_1.4 (*top*) or Ca_v_1.1 (*bottom*). NALCN are domain colored and Na_v_1.4 and Ca_v_1.1 are colored in slate and light pink, respectively. An intracellular view and a side view are shown for each comparison. The major conformational changes from Na_v_1.4 or Ca_v_1.1 to NALCN are indicated by red arrows.

It is intriguing that the position of the III-IV linker in NALCN is distinct from those of both Na_v_1.4 and Ca_v_1.1, in different ways. NALCN does not have an IFM motif, a highly conserved motif in the III-IV linker of all Na_v_ channels that is crucial for fast inactivation through an allosteric inhibition mechanism (Armstrong and Bezanilla, 1977; Yan et al., 2017). In Na_v_1.4, the III-IV linker is away from the inner pore, and its helix rotates counterclockwise by about 75^0^ but at a similar vertical height compared to that of NALCN (Figure 6C, *top*). The helix in the III-IV linker, however, shows a similar orientation but distinct vertical height in Ca_v_1.1. Compared to Ca_v_1.1, the helix in the III-IV linker of NALCN is about 6 Å up shifted towards the extracellular side, probably due to a shorter loop connecting S6_III_ and the helix in the III-IV linker (Figure 6C, *bottom*). The II-III linker, which is part of the elongated S6_II_ in Ca_v_1.1, bends towards the center pore. These structural features in NALCN make S6_I_, II-III linker and III-IV linker form a close contact that tightly seal the inner gate.

Notably, S5_II_ and S6_II_ are much closer to the center pore in NALCN compared with Na_v_1.4/Ca_v_1.1. The adjacent S4-5_II_ also displays significant conformational changes in NALCN (Figure 6C). The dramatic differences in the pore domain of repeat II are closely related to the binding of a lipid molecule (Figure 2C). The lipid, only observed in NALNC, may play an important role in regulating the gating of NALCN.

## Discussion

Our study reports the high-resolution structure of NALCN in complex with the auxiliary subunit FAM155A. The structure reveals unique features of NALCN that help explain its ion selectivity, voltage sensing and specific interaction with auxiliary subunits. It also provides an important framework for comparative investigations of channel properties of related Na_v_/Ca_v_ channels.

Unlike Na_v_/Ca_v_ channels, there is only one subtype of NALCN in most organisms including human. Severe mutations/deletions of NALCN can be fatal as there are no channels with redundant functions. This idea is supported by the evidence that NALCN is essential for neonatal survival and NALCN knockout mice die within 24 hours after birth (Lu et al., 2007). We summarized the reported disease-related mutations and mapped them onto the structure (Figure S11A, Table S3). We found that most of the mutations are located on the pore domain of NALCN (Figure S11A). These mutations may affect NALCN function by directly altering ion transduction properties through the pore. Two mutations on the III-IV linker, T1165P and R1181Q, may lead to changes of channel gating through affecting the local interactions of the III-IV linker with adjacent S6_I_, the II-III linker, S6_IV_ and CTD. Our structure thus provides an important framework to interpret the potential disease-causing mechanisms of the reported mutations.

TTX is a neurotoxin and of sodium channel blocker. The Na^+^ current conducted by NALCN is TTX-resistant (Lu et al., 2007). Our structure immediately illustrates the incompatibility of TTX-bound in the pore of NALCN. Previous structural comparison between the TTX bound Na_v_1.7 and apo Na_v_1.5 revealed that a residue with an aromatic ring (Tyr362 in Na_v_1.7) in repeat I is the structural determinant for TTX sensitivity in Na_v_ channels (Jiang et al., 2020). This residue forms a π-π stacking interaction with TTX bound in the outer pore of Na_v_1.7 and replacement of the residue with a cystine results in much reduced TTX binding affinity in TTX-insensitive Na_v_1.5. The overall backbone of the pore helices and the other residues that mediate the interaction with TTX are almost the same between Na_v_1.7 and Na_v_1.5, though (Figure S11B). In NALCN, not only the determinant residue is replaced by a glycine, but the other key residues that involved in TTX binding are also not conserved (Figure S8). Moreover, the overall backbone of the pore helices has considerably changed compared with Na_v_1.7, which lead to clashes with the potential TTX binding site (Figure S11B). These structural features of NALCN make it unable to bind TTX, even more resistant to TTX than the TTX-insensitive Na_v_ channels.

The relative GCs positions in NALCN VSDs are the same as that of Na_v_1.4 and Ca_v_1.1, except for the R5 in VSDIII where the side chain of arginine residue faces below the occluding residue in S2. However, the Cα atom of R5 is not shifted, indicating all the VSDs in NALCN are in up conformations (Figure S10). Further comparisons with Na_v_PaS whose VSDIII and VSDIV are in a potential resting state revealed obvious upward movement of the S4 segments in VSDIII and VSDIV from Na_v_PaS to NALCN, confirming the depolarized conformations of VSDs in NALCN (Figure S10). Although our structure seems to be in a similar state with the inactivated Na_v_/Ca_v_ structures (Shen et al., 2019; Wu et al., 2016), the exact state of NALCN channel in our structure remains to be further investigated considering its unique properties.

The auxiliary subunits play an important role in fine tuning the activity of NALCN. FAM155A, previously characterized as an ER-resident protein in *C. elegans* (Xie et al., 2013), turns out to interact with NALCN directly in humans. It may help NALCN to fold stably and translocate to cell membrane. Our observation of an obvious left shift of the G-V curve when UNC79 and UNC80 were co-expressed with NALCN and FAM155A has also suggested a regulatory effect of UNC79 and UNC80 on NALCN. NALCN contains a π helix in each S6 segment (Figure 5A). Transition of the π helix to regular α helix during channel conformational changes have been reported in many structures (McGoldrick et al., 2018; Singh et al., 2018; Su et al., 2018; Zhao et al., 2019). Association of UNC79, UNC80 into NALCN-FAM155A subcomplex may induce conformational changes and opening of the pore through π-to-α helical transition in S6. Additional components, such as several GPCRs and SFKs may also contribute to the modulation of NALCN function. It remains an open question whether there are more auxiliary subunits of the NALCN channelosome. Future structural and functional studies on larger complexes of NALCN will deepen our understanding of the regulation of the NALCN channelosome.

## Acknowledgements

We thank the Cryo-EM Facility of Westlake University for providing cryo-EM and computation support. We thank Westlake University Supercomputer Center for computational resource and related assistance. We thank the Mass Spectrometry & Metabolomics Core Facility at the Center for Biomedical Research Core Facilities of Westlake University for MS sample analysis. We thank Dr. Stephan Pless (University of Copenhagen) for kindly sharing the plasmids of human UNC-79, UNC-80 and FAM155A. We thank Dr. Hongtao Yu (Westlake University), Dr. Nieng Yan (Princeton University) for critical reading of the manuscript. This work was supported by Westlake Education Foundation.

## Author Contributions

J.W. conceived the project. J.W., Z.Y., F.Y. designed all experiments and supervised the project. J.X., and Z.Y. prepared the protein sample, J.X., M.K. and J.W. collected cryo-EM data and calculated the cryo-EM map. J.W. built the atomic model. M.K. performed the MD simulations. L.X. conducted the electrophysiological experiments. All authors contributed to data analysis. J.W. and Z.Y. wrote the manuscript.

## Supplementary Figure Legends

**Figure S1. Electrophysiological properties of NALCN characterized by patch-clamp recordings, Related to Figures 1–4.**

**(A)** Representative whole-cell recordings of HEK293 with mock transfection and NALCN channel transfected, respectively. Dashed line indicates the zero-current level. **(B)** Representative whole-cell recordings of NALCN, FAM155A, UNC79 and UNC80 co-expressed in HEK293 cells. Data points in shaded rectangle were averaged for plotting current-voltage curve in **(D)**. Dashed line in black indicates the zero-current level. Dashed curves in red and blue are fitted curves to the inactivation and activation kinetics with exponential functions, respectively. NALCN current was inhibited by 1 mM verapamil and the current could be recovery after washing off. The same whole-cell patch was used during verapamil inhibition. **(C)** Representative whole-cell recordings of NALCN channel and FAM155A co-expressed in HEK293 cells. Data points in shaded rectangle were averaged for plotting current-voltage curve. **(D)** Current-voltage curve of NALCN co-expressed with FAM155A only (n = 6) and co-expressed with FAM155A, UNC79 and UNC80 (n = 7). The currents were normalized to the maximum current level. **(E)** Conductance-voltage (G-V) curves of NALCN co-expressed with FAM155A (Red: V_1/2_ = −44.66 ± 1.91 mV; apparent gating charge (Z_app_) = 1.22 ± 0.18 e_0_) and co-expressed with FAM155A, UNC79 and UNC80 (Blue: V_1/2_ = −65.07 ± 2.11 mV; apparent gating charge (Z_app_) = 1.25 ± 0.55 e_0_). G-V curves were fitted to a single-Boltzmann function (equation (1) in Methods, dash curves). Co-expression with FAM155A, UNC79 and UNC80 shifted the G-V curve of NALCN channel towards the hyperpolarization voltage. All data are given as mean ± s.e.m. **(F)** Representative whole-cell recordings of the steady-state inactivation of NALCN co-expressed with FAM155A (*left*) or with FAM155A, UNC79 and UNC80 (*right*). **(G)** Prepulse voltage dependence of steady-state inactivation of NALCN channel co-expressed with auxiliary subunits measured from −80 mV (n = 7).

**Figure S2. Cryo-EM analysis of human NALCN in complex with FAM155A, Related to Figure 1 and Table S1.**

**(A)** Last step purification of recombinantly expressed NALCN-FAM155A complex. A representative chromatogram of gel filtration purification is shown. The indicated fractions were resolved by SDS-polyacrylamide gel electrophoresis followed by Coomassie blue staining. The bands corresponding to NALCN and FAM155A are indicated by black and orange arrows, respectively. The protein identity of each band was confirmed by mass spectrometric analysis. **(B)** Representative micrograph of the NALCN-FAM155A complex. Scale bar, 40 nm. **(C)** Representative two-dimensional class averages for NALCN-FAM155A complex. Box size: 240 Å; circle mask: 220 Å. **(D)** Angular distribution of the particles of the final reconstruction generated by cryoSPARC. **(E)** Gold standard FSC curves for the 3D reconstructions. The curves for the reconstructions of the overall map, map of the intracellular region and map of the extracellular region are indicated by blue, yellow, and red lines, respectively. **(E)** Validation of the final structure models. FSC curves of the final refined model versus the corresponding map that it was refined against (black); of the model refined in the first of the two independent half maps used for the gold-standard Fourier shell correlation curves versus that same map (blue); and of the model refined in the first of the two independent maps versus the second independent map (red). The small difference between the blue and red curves indicates that the refinement of the atomic coordinates did not suffer from overfitting.

**Figure S3. Flowchart for EM data processing, Related to Figure 1 and Table S1.**

Please refer to “Image processing” session in Methods for details.

**Figure S4. EM maps of human NALCN, Related to Figures 1–6.**

**(A)** EM maps for the S1-S6 segments in the four repeats of NALCN. The side groups of representative bulky residues are labeled. The gating charges in S4 helices are highlighted in brown. The densities, shown as blue meshes, are contoured at 4-5 σ in PyMol. **(B)** EM maps for the two resolved lipid molecules that located in the inner side of the membrane. Two perpendicular views are shown for each lipid. The lipid densities are contoured at 3-4 σ in PyMol.

**Figure S5. EM maps of NALCN-FAM155A complex, Related to Figures 1–5.**

**(A)** All residues that constitute the SF in four repeats of NALCN are clearly resolved. The densities are contoured at 4-5 σ in PyMol. The EEKE residues are labeled orange. **(B)** Representative EM maps of FAM155A. The side groups of bulky residues are labeled. **(C)** EM maps of the disulfide bonds in the complex structure. The densities are contoured at 3-4 σ in PyMol. **(D)** EM maps of the sugar moieties in the glycosylation sites. The densities are contoured at 2-4 σ in PyMol.

**Figure S6. Mass Spectrometric analysis of crosslinked NALCN-FAM155A complex, related to Figure 3.**

Shown here is a high resolution HCD spectra of the inter-subunit crosslinked peptides from NALCN-FAM155A complex. The number shown in the brackets refer to the residue number in the indicated subunit. Inset: Structure of the NALCN-FAM155A interaction interface. K1377 of NALCN is shown in stick. The C-terminal residue in the resolved FAM155A structure is E382, which is about 27 Å from K1377 of NALCN. K395 of FAM155A, 13 residues after E382, is likely to be in the vicinity of cross-linking range with K1377 of NALCN. The cross-linking MS result is consistent with our structure observation.

**Figure S7. Interaction interfaces between NALCN and FAM155A, Related to Figure 3 and Table S2.**

**(A)** Structure of the FAM155A. The N-lobe and C-lobe were colored in gold and orange, respectively. The secondary elements are labeled based on the definition in Figure 3. **(B)** Crystal structure of the CRD domain of MuSK and its superimposition with FAM155A. The conserved disulfide bonds between the two structures are highlighted by black arrows. **(C)** Electrostatic surface of the interaction interfaces between NALCN and FAM155A. The structures are shown with both subunits in electrostatic surface (left), with NALCN in cartoon and FAM155A in electrostatic surface (middle), or with FAM155A in cartoon and NALCN in electrostatic surface (right). Two side views are presented. The green circles highlight the interaction interfaces that have complementary electrical potential between NALCN and FAM155A.

**Figure S8. Sequence alignment among NALCN and selected Na_v_/Ca_v_ channels, Related to Figure 3–6 and Table S2, S3.**

The primary sequences of four human Na_v_ subtypes and three human Ca_v_ subtypes are compared with that of human NALCN using Clustal W (Larkin et al., 2007). The invariant residues are shaded gray and the conserved residues are colored red. Secondary structural elements of human NALCN are presented above the sequence alignment and color-coded for each repeat. The critical EEKE residues in the selectivity filter are colored red and shaded yellow. The gating charge residues (labeled R1-R6) in each S4 segments are colored white and shaded red. The residues on S1-S3 that may facilitate the gating charge transfer, including the labeled An1 and An2 sites and the occluding residues on S2, are shaded cyan. The residues that form disulfide bonds are labeled with squares below the sequence alignment. The glycosylation sites are labeled with black triangles and the residues that mediate interaction with FAM155A are labeled with orange triangles. The Uniprot IDs for the aligned sequences are: hNALCN: Q8IZF0; hNav1.2:Q99250; hNa_v_1.4:P35499; hNa_v_1.5:Q14524; hNa_v_1.7:Q15858; hCa_v_1.1: Q13698; hCa_v_2.1: O00555; hCa_v_3.1: O43497.

**Figure S9. Sequence alignment of FAM155A, Related to Figure 3 and Table S2.**

The primary sequences of FAM155A from human, mouse, and bovine and FAM155B from human are aligned using Clustal W. The invariant residues are colored white and shaded gray. The conserved residues are colored orange. The secondary structure elements of human FAM155A are indicated above the sequence alignment. The residues that form disulfide bonds are labeled with squares below the sequence alignment. The glycosylation site is labeled with black triangle and the residues that mediate interaction with NALCN are labeled with orange triangles. The Uniprot IDs for the aligned sequences are: hFAM155A: B1AL88, mFAM155A: Q8CCS2; bFAM155A: A4IFM1, hFMA155B:O75949.

**Figure S10. All four VSDs adopt up conformations, Related to Figure 4.**

Each VSD of NALCN is superimposed with their counterpart in Na_v_1.4 (PDB: 6AGF) and Ca_v_1.1 (6JP5). The VSDs are aligned relative to CTC residues and An1 on S2 (labeled blue). S1 segments were omitted by visual clarity. The gating charges (GCs), charge transfer center (CTC) and An1 on S2 are shown as stick and Cα atoms of the GCs are shown as spheres. The GCs on S4 are labeled as R1-R6. The positions of the GCs in NALCN are similar as that of Na_v_1.4 and Ca_v_1.1, which represent up or depolarized conformations. By contrast, the positions of GCs in VSDIII and VSDIV show an upward shift compared to that of Na_v_PaS, whose VSDIII/IV may represent resting state.

**Figure S11. Disease mutations mapping on NALCN and TTX incompatibility of NALCN, related to Figure 5 and Table S3.**

**(A)** The structure of NALCN in cartoon are shown in both side and extracellular views. The Cα atoms of the residues whose mutations lead to human diseases are shown as spheres. **(B)** Superimpositions of the pore helices and selectivity filter of NALCN with that of Na_v_1.7 bound with TTX (*left*). The determinant Tyr362 in Na_v_1.7 is not conserved in NALCN. The structure differences in the pore region of NALCN result in its potential clashes with the TTX binding site. As a comparison, the TTX-insensitive Na_v_1.5 does not process the determinant Tyr but show a similar backbone structure with TTX-bound Na_v_1.7 (*right*).

## METHOD DETAILS

### Protein expression and purification

NALCN, UNC-80, UNC-79 and FAM155A were cloned into pCAG vector. HEK 293F cells (Invitrogen) were transfected with the four plasmids and harvested after 60 hours. Cells were resuspended in buffer containing 25 mM Mops (pH 7.4), 250 mM NaCl, 0.5% (w/v) Lauryl Maltose Neopentyl Glycol (LMNG, Anatrace), 0.06% (w/v) cholesteryl hemisuccinate Tris salt (CHS, Anatrace) and protease inhibitor cocktail including 2 mM phenylmethylsulfonyl fluoride (PMSF), 1.3 μg/ml aprotinin, 0.7 μg/ml pepstatin, and 5 μg/ml leupeptin, then incubated at 4°C for 2 hours. The insoluble fraction was precipitated by ultracentrifugation at 255,700 g for 1 h, and the supernatant was applied to anti-Flag M2 affinity resin (GenScript) by gravity. The resin was rinsed several times with the Wash buffer, which contains 25 mM Mops (pH 7.4), 500 mM NaCl, 0.01% (w/v) Lauryl Maltose Neopentyl Glycol, and the protease inhibitor cocktail. The target proteins were eluted by the Elution buffer containing 25 mM Mops (pH 7.4), 500 mM NaCl., 0.02% (w/v) glyco diosgenin (GDN, Anatrace), and the protease inhibitor cocktail plus 200 μg/ml FLAG peptide (GenScript). The eluent was then concentrated using a 50-kDa cut-off Centricon (Millipore) and further purified by Superose 6 Increase 10/300 GL column (GE Healthcare) in 25 mM Mops (pH 7.4) 150 mM NaCl, 0.02% (w/v) GDN, and the protease inhibitor cocktail. Fractions were collected and concentrated to about 8.5 mg/ ml for cryo-EM analysis.

### Whole-cell electrophysiology

Patch-clamp recordings were performed with a HEKA EPC10 amplifier with PatchMaster software (HEKA) in whole-cell configuration. Patch pipettes were prepared from borosilicate glass and fire-polished to resistance of ~ 6 MΩ. For whole cell recording, serial resistance was compensated by at least 65%. Extracellular solution contained 150 mM NaCl, 10 mM Hepes, and 30 mM d-(+)-glucose (pH 7.4) with NaOH, and intracellular solution contained 136 mM NaCl, 5 mM EGTA, 10 mM Hepes, and 2 mM Na2ATP (adenosine 5′-triphosphate) (pH 7.2) with NaOH. Current amplitude at the steady state during the last 20 ms of voltage steps was averaged to construct the G-V curve. All recordings were performed under room temperature (~22°C). Temperature variation was less than 1 °C as monitored by a thermometer. Current signal was sampled at 20 kHz.

HEK-293T cells were grown in Dulbecco’s modified Eagle’s medium (Thermo Fisher Scientific) supplemented with 10% fetal bovine serum and 1% penicillin-streptomycin at 37°C in a 5% CO2 humidified growth incubator. Cells were transiently transfected by Lipofectamine 3000 (Life Technologies) 48-72 hours before patch-clamp recordings. The NALCN, UNC-79, UNC-80, and FAM155A cDNAs were mixed in the mass ratio of 1:1:1:1.

To apply solutions containing ligands such as verapamil during patch-clamp recording, a rapid solution changer with a gravity-driven perfusion system was used (RSC-200, Bio-Logic). Each solution was delivered through a separate tube so that there was no mixing of solutions. Pipette tip with a membrane patch was placed directly in front of the perfusion outlet during recording. Each membrane patch was recorded for only once.

Data from patch-clamp recordings were analyzed in Igor Pro (WaveMatrix). To characterize the steady-state G-V curves, a single-Boltzmann function was used:

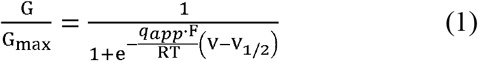

where G/G_max_ is the normalized conductance, V_1/2_ is the half-activation voltage, q_app_ is the apparent gating charge and F is Faraday’s constant, R is gas constant, and T is temperature in Kelvin, which is set to 295 K (22°C).

### Cryo-EM sample preparation and data collection

The purified NALCN complex was concentrated to about 8.5 mg/ml for cryo-sample preparation. Aliquots (3.5 μl) of protein solution were loaded onto glow-discharged holey carbon grids (Quantifoil Au R1.2/1.3), which were blotted for 3.5 s and immersed in liquid ethane cooled by liquid nitrogen using Vitrobot (Mark IV, Thermo Fisher Scientific). The grids were exposed through Titan Krios operating at 300 kV equipped with Gatan K3 Summit detector and GIF Quantum energy filter in super-resolution mode (81,000× magnification). Movie stacks were automatically acquired using AutoEMation (Lei et al., 2005), with a 20 eV slit width and a defocus range from −0.5 μm to −2.5 μm. Each stack consisting of 32 frames was exposed for 2.56 s with 0.08 s per frame, and for approximately 50 e^-^/Å^2^ of total dose.

### Cryo-EM data processing

Collected movie stacks were motion-corrected by MotionCor2 (Zheng et al., 2017) with 2-fold binning, producing micrographs of 1.087 Å pixel size. Following patch-CTF estimation, around eleven million particles were automatically picked with cryoSPARC v2 (Punjani et al., 2017) from 17,922 micrographs. Three rounds of 2D classification enriched images of good classes, resulting in a total of 905,458 particles being selected. An Ab-initio reconstruction generated a low-resolution initial map for the subsequent homogeneous and non-uniform refinement jobs (Ali Punjani, 2019), yielding a 3.8-Å reconstruction with identifiable side-chain densities in NALCN pore domain only. To improve map quality, we exported the aligned particles and did a 3D classification without orientation assignment using RELION 3.0 (--skip-align flag) (Scheres, 2012; Zivanov et al., 2018). One out of five 3D classes represented clear secondary structures of the overall channel complex, and the 65,177 particles were fed back to cryoSPARC v2 for further homogeneous and non-uniform refinements that pushed the resolution to 3.14 Å. Adapted masks for protein regions, extracellular subunit, and intracellular domain were applied in particle subtraction and local refinement tasks, which helped in achieving slightly higher resolution at 3.11 Å, 2.94 Å and 3.04 Å, respectively. Map resolutions were determined by the gold-standard Fourier shell correlation (FSC) 0.143 criterion using Phenix.mtriage (Adams et al., 2010).

### Model building and refinement

An initial model of NALCN was generated using SWISS-MODEL online server (Waterhouse et al., 2018). The template used for the homology modeling was the α1 subunit of Ca_v_1.1 (PDB: 5GJV). The model was firstly docked into the final reconstruction map in Chimera (Pettersen et al., 2004) and then manually adjusted in COOT (Emsley and Cowtan, 2004). After building the model of NALCN, an extra density remained unmodelled which is close to the ECL of NALCN in the extracellular side. The high quality of map enables us to identify that the extra density belongs to FAM155A. The structure of FAM155A was then *de novo* built. Sequence assignment was guided by bulky residues such as Phe, Tyr and Trp. Up to six disulfide bonds and a glycosylation site (Asn217) were identified in FAM155A, further verifying the accuracy of the model. Additional lipid and detergent molecules were manually built to fit into the corresponding densities. In total, 1295 and 182 residues were built for NALCN and FAM155A respectively. The side chains of most of the built residues were assigned except for a short α helix at the C-terminus of NALCN. This short helixis disconnected to any other part of the structure while it is in close contact with the CTD. We tentatively assigned it as part of the CTD (1688-1704) based on sequence and local density analysis. However, there is a possibility that this helix may belong to other interaction proteins such as UNC80, which are not resolved in our current structure.

Subsequently, the models were refined against the corresponding maps by PHENIX (Adams et al., 2010) in real space (phenix.real_space_refine) with secondary structure and geometry restraints generated by ProSMART (Nicholls et al., 2014). Overfitting of the overall models was monitored by refining the models in one of the two independent half maps from the gold-standard refinement approach and testing the refined model against the other map (Amunts et al., 2014). Statistics of 3D reconstruction and model refinement can be found in Table S2.

### Molecular dynamics simulation

The pore domain of NALCN (residue 171-204, 260-328, 490-603, 1001-1042, 1094-1163, 1318-1400, and 1420-1455) was embedded in 1-palmitoyl-2-oleoyl-sn-glycero-3-phosphocholine (POPC) bilayer for molecular dynamics simulations. After determination of side-chain protonation states by PROPKA3.1 (Olsson et al., 2011) and peptide terminal neutralization by acetyl/methylamine capping groups, water molecules and 150 mM NaCl were added using VMD (Humphrey et al., 1996), resulting in a system of size 100□×□100□×□105 Å with ~93,000 atoms. The system was parameterized using tleap in AmberTools18 by ff14SB (Maier et al., 2015) and LIPID17 force fields, and along with TIP3P water model. All the simulations were performed using OpenMM7 (Eastman et al., 2017). We first applied positional constraints (k=10 kcal/mol/Å^2^) on protein heavy atoms for a 5000-step minimization, then released constraints of side-chains when heating up the system to 310 K with 2 fs stepsize (H-bonds constraints), 1 ps-1 friction coefficient for Langevin dynamics, Particle Mesh Ewald (PME) method, and 12 Å non-bonded cutoff. During 50-ns pre-equilibration for membrane relaxation in NPT ensemble, the force constant k of backbone atoms was reduced from 10 to 0.1 kcal/mol/Å^2^. Three independent 200-ns production runs were conducted constraining Cα atoms more than 15 Å away from the selectivity filter residues (EEKE) with k = 0.1 kcal/mol/Å^2^. Simulation trajectories were analyzed using MDTraj 1.9.3 (McGibbon et al., 2015). After superimposition of the pore region to the initial structure, we recorded the positions of sodium ions within 15 Å of SF residues EEKE every 1 ns for each run.

### Protein crosslinking and LC-MS/MS analysis

Equal amount (w/w) of BS3 (bis[sulfosuccinimidyl] suberate) was added into the protein mixture, which was incubated at room temperature for 1 h. The reaction was terminated by adding 0.5 M ammonium bicarbonate to a final concentration of 20 mM for 10 min incubation. The SDS-PAGE was used to separate the crosslinked protein and stained with Coomassie Blue G-250. The gel bands of interest were cut into pieces. Sample was digested by trypsin with prior reduction and alkylation in 50 mM ammonium bicarbonate at 37°C overnight. The digested products were extracted twice with 1% formic acid in 50% acetonitrile aqueous solution and dried to reduce volume by speedvac.

For LC-MS/MS analysis, the peptides were separated by a 65 min gradient elution at a flow rate 0.300 μL/min with the Thermo EASY-nLC1200 integrated nano-HPLC system which is directly interfaced with the Thermo Q Exactive HF-X mass spectrometer. The analytical column was a home-made fused silica capillary column (75 μm ID, 150 mm length; Upchurch, Oak Harbor, WA) packed with C-18 resin (300 A, 3 μm, Varian, Lexington, MA). Mobile phase A consisted of 0.1% formic acid, and mobile phase B consisted of 100% acetonitrile and 0.1% formic acid. The mass spectrometer was operated in the data-dependent acquisition mode using the Xcalibur 4.1 software and there is a single full-scan mass spectrum in the Orbitrap (300-1800 m/z, 60,000 resolution) followed by 20 data-dependent MS/MS scans at 30% normalized collision energy. Each mass spectrum was analyzed using the Thermo Xcalibur Qual Browser and pLink 2 (Chen et al., 2019) for the database searching and crosslinking analysis.

## DATA AND SOFTWARE AVAILABILITY

### Data Resources

Atomic coordinate and corresponding EM maps of the NALCN-FAM155A complex (PDB: 7CM3; EMDB: EMD-30400) have been deposited in the Protein Data Bank (http://www.rcsb.org) and the Electron Microscopy Data Bank (https://www.ebi.ac.uk/pdbe/emdb/).

### Supplementary Movies

**Movie S1. MD simulation of NALCN pore domain for sodium binding site identification, Related to Figure 5.**

Left panel. Top view of selectivity filter. Ion binding residues are highlighted in stick representation. Probability density of sodium is illustrated in mesh grid. Sodium ions are colored in magenta. Right panel. Side views of selectivity filter.

### Supplementary Tables

**Table S1. Data collection and model statistics, Related to Figure 1.**

**Table S2. Summary of contact interfaces between NALCN and FAM155A, Related to Figure 3.**

**Table S3. Disease related mutations on human NALCN, related to Figure S12.**

