## Supplementary figures and images for "Structure of human sodium leak channel NALCN in complex with FAM155A"

### Figure S1

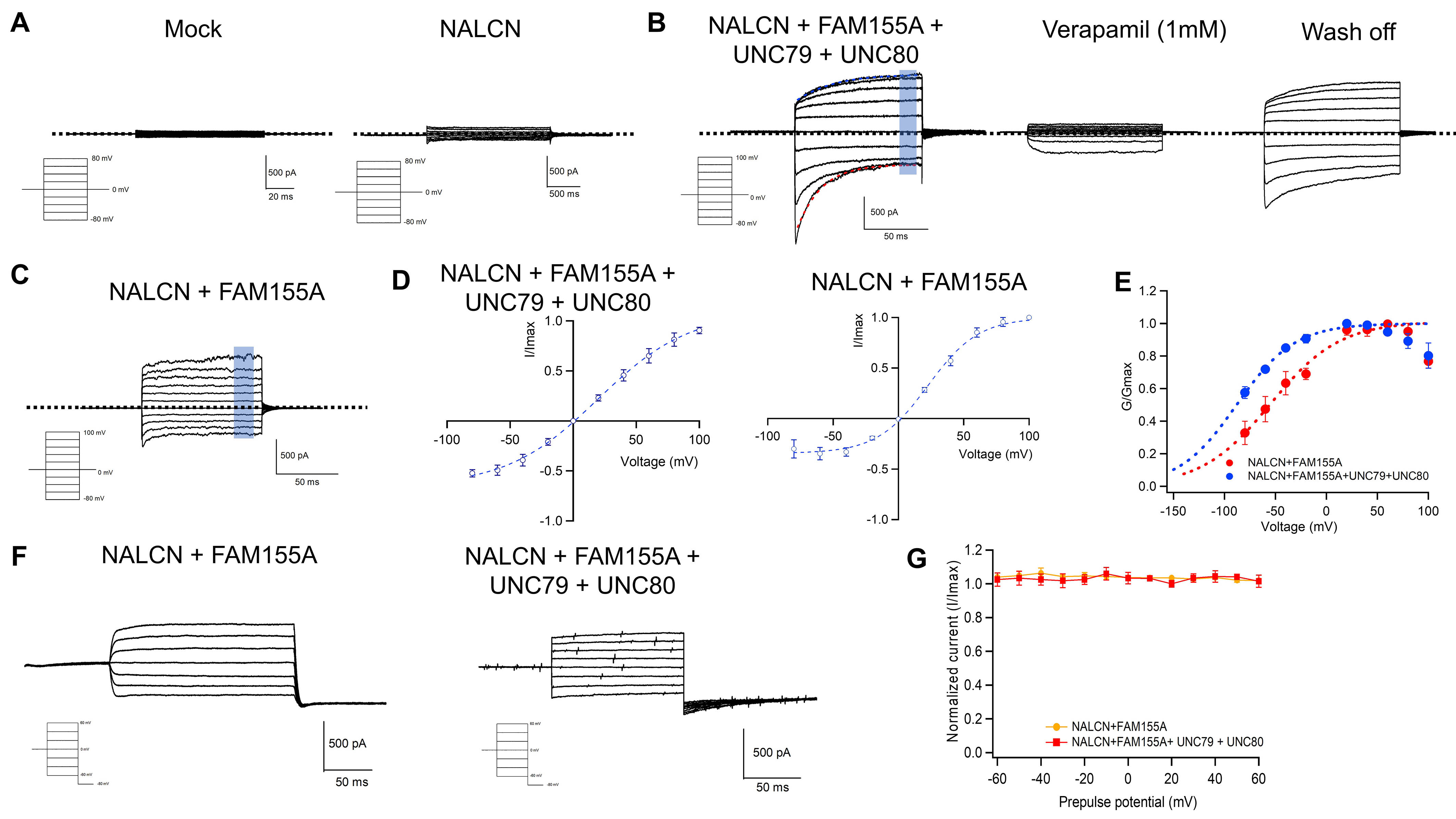

### Figure S2

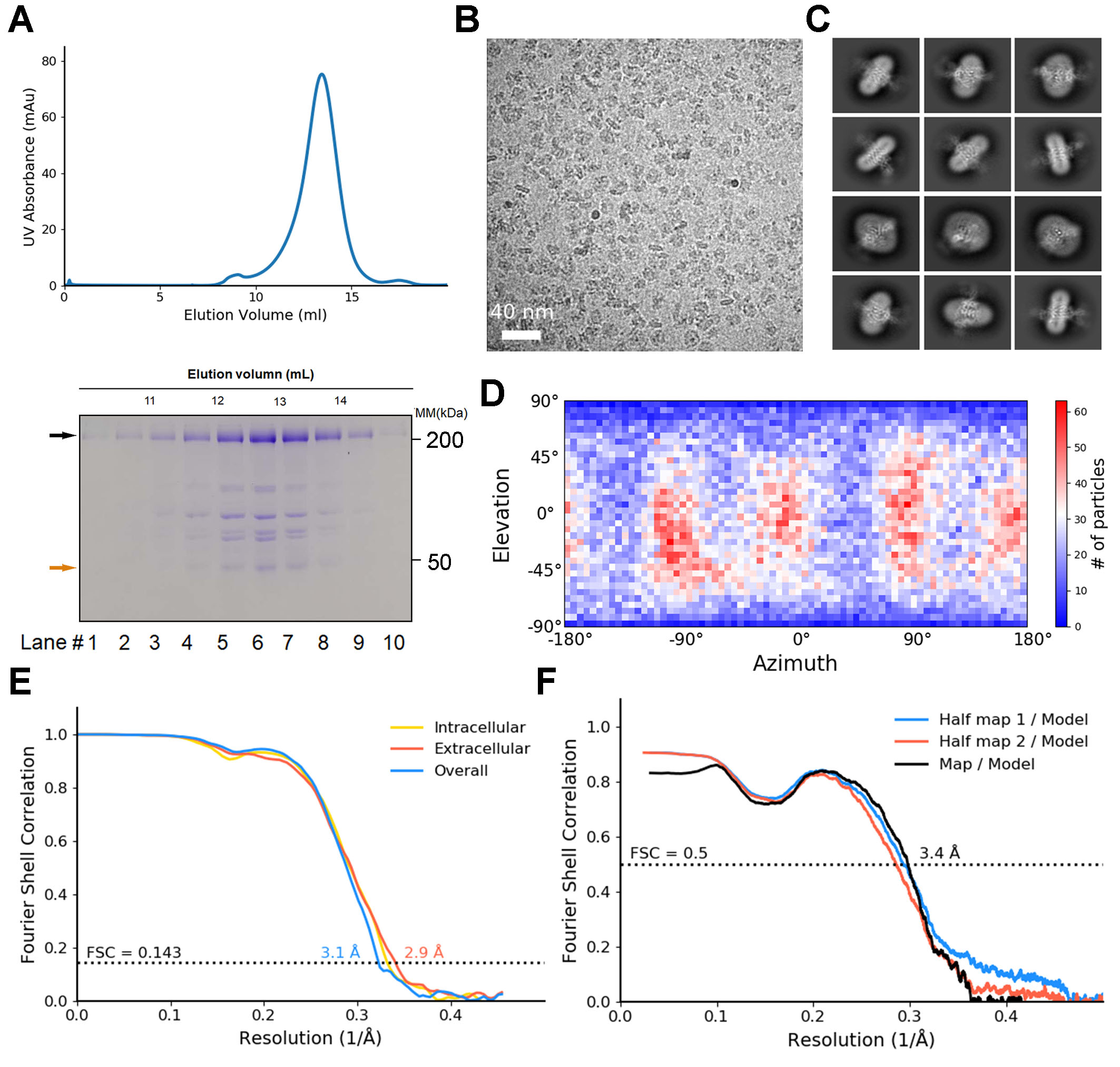

### Figure S3

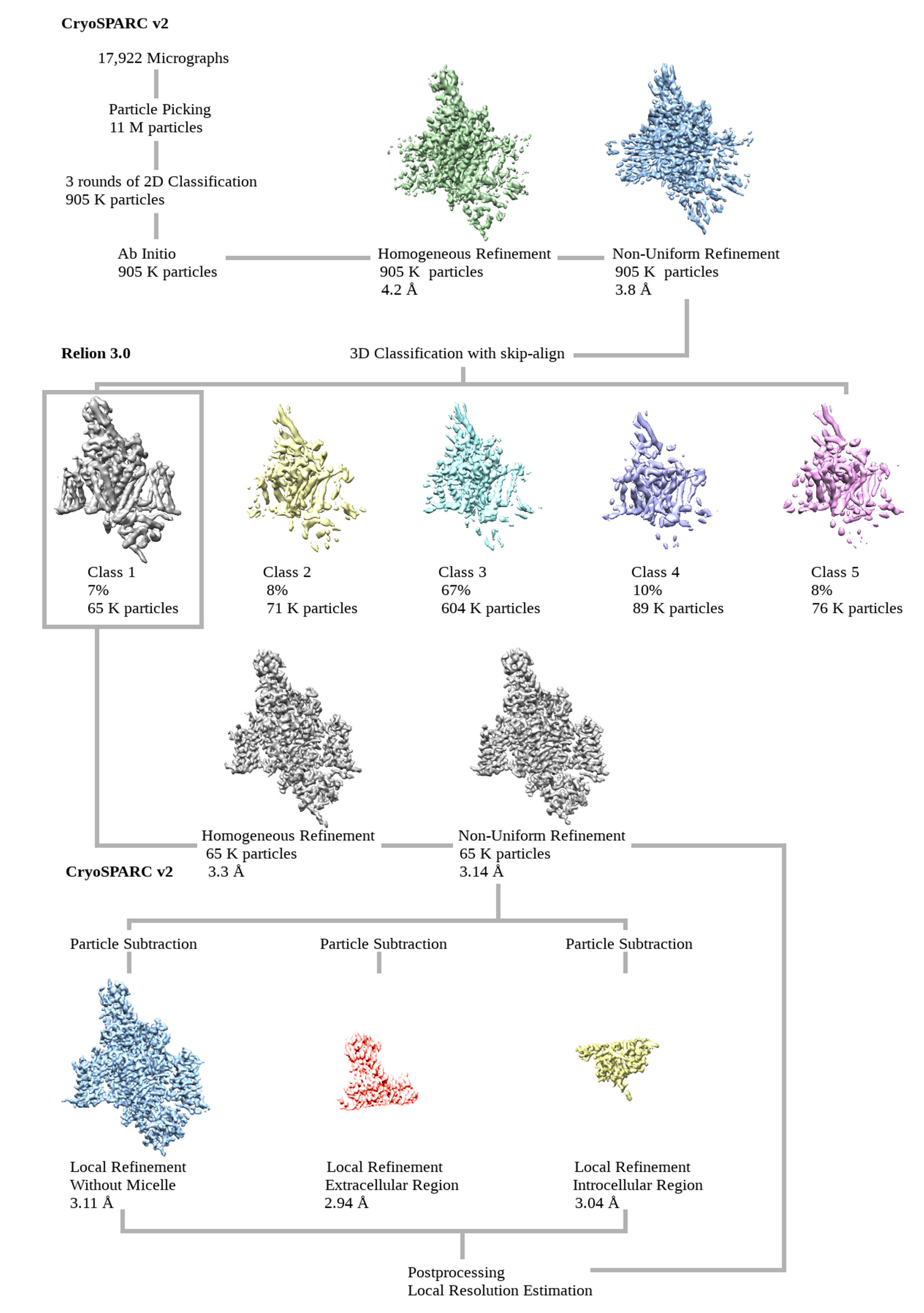

### Figure S4

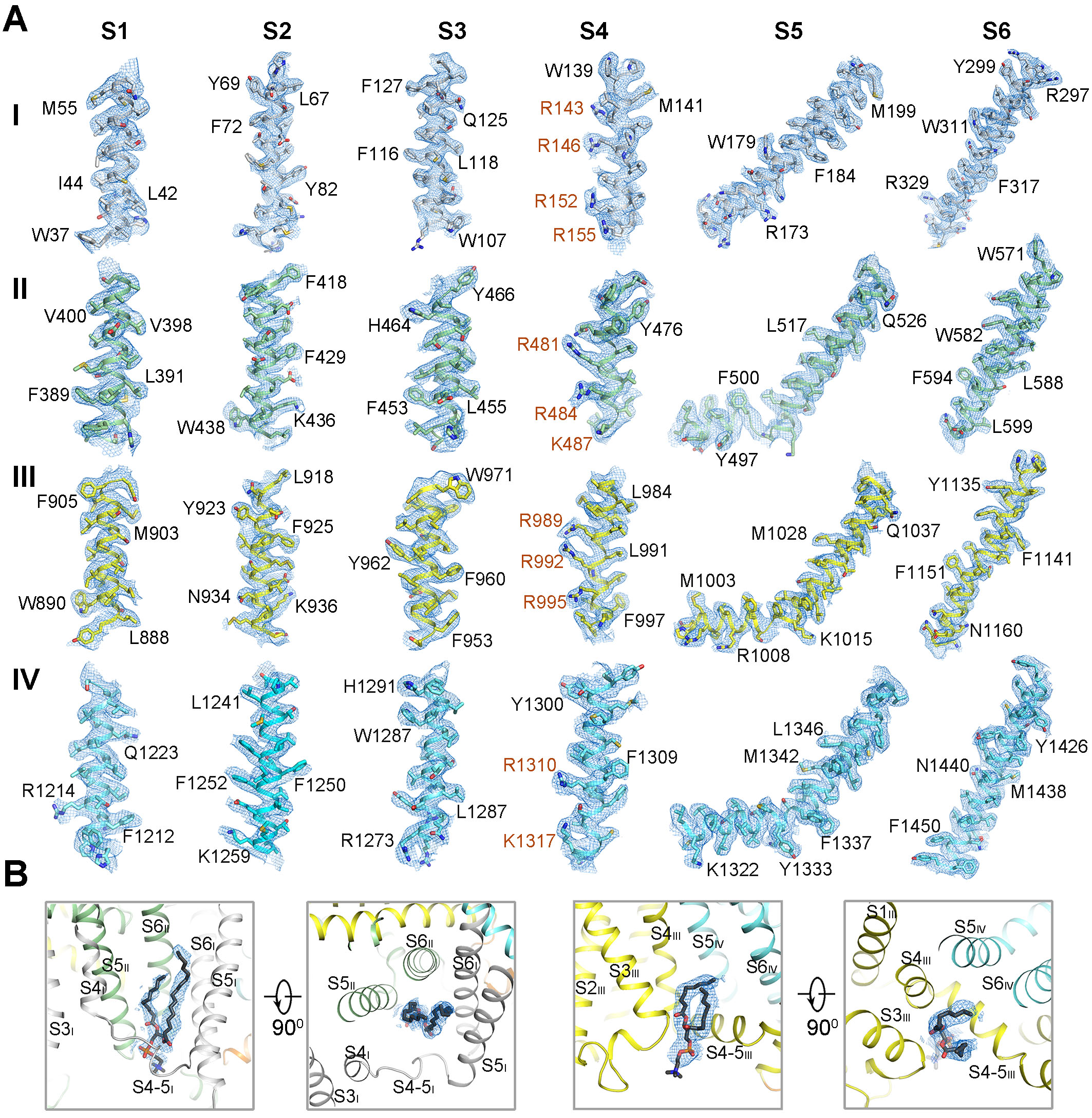

### Figure S5

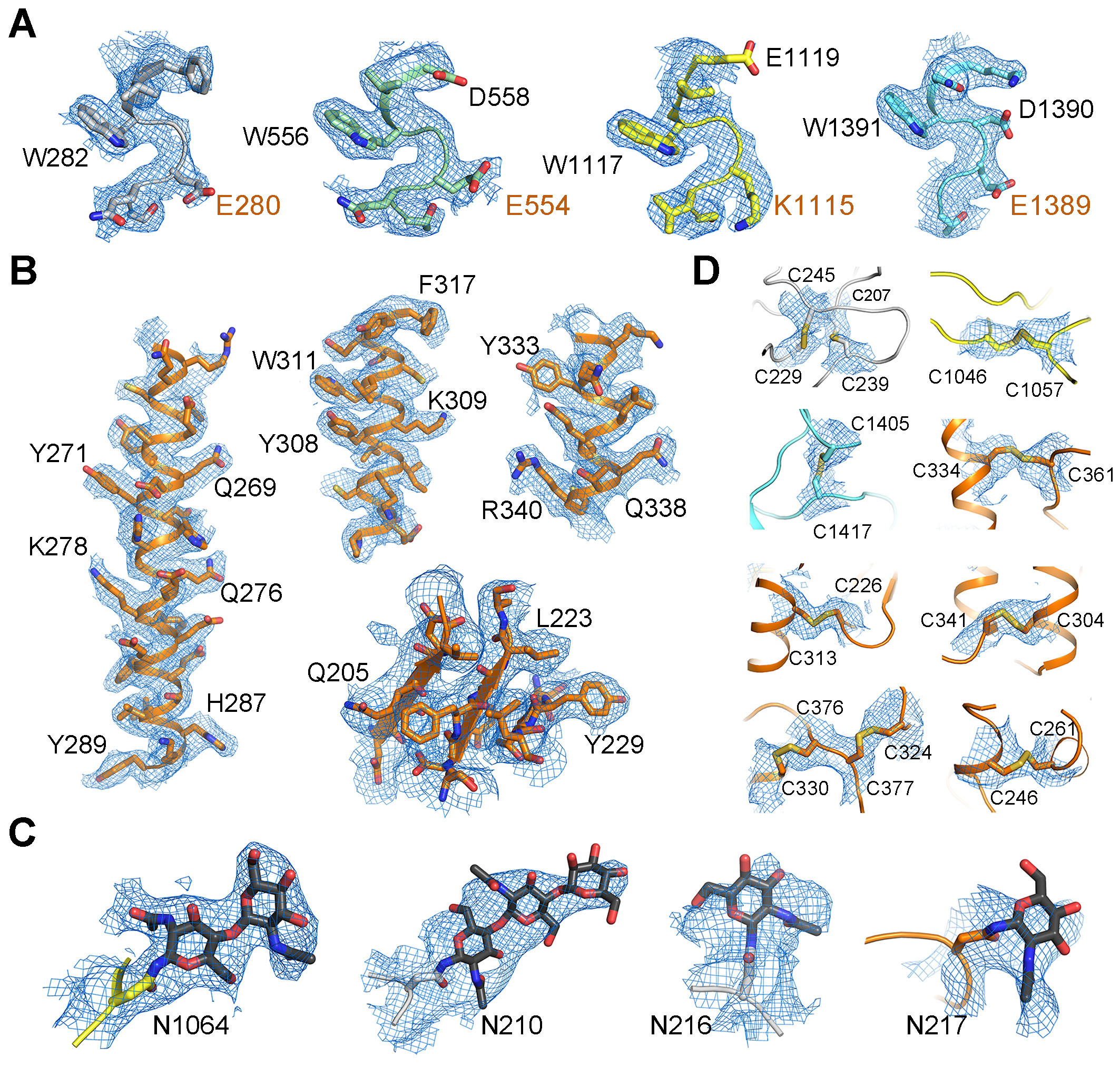

### Figure S6

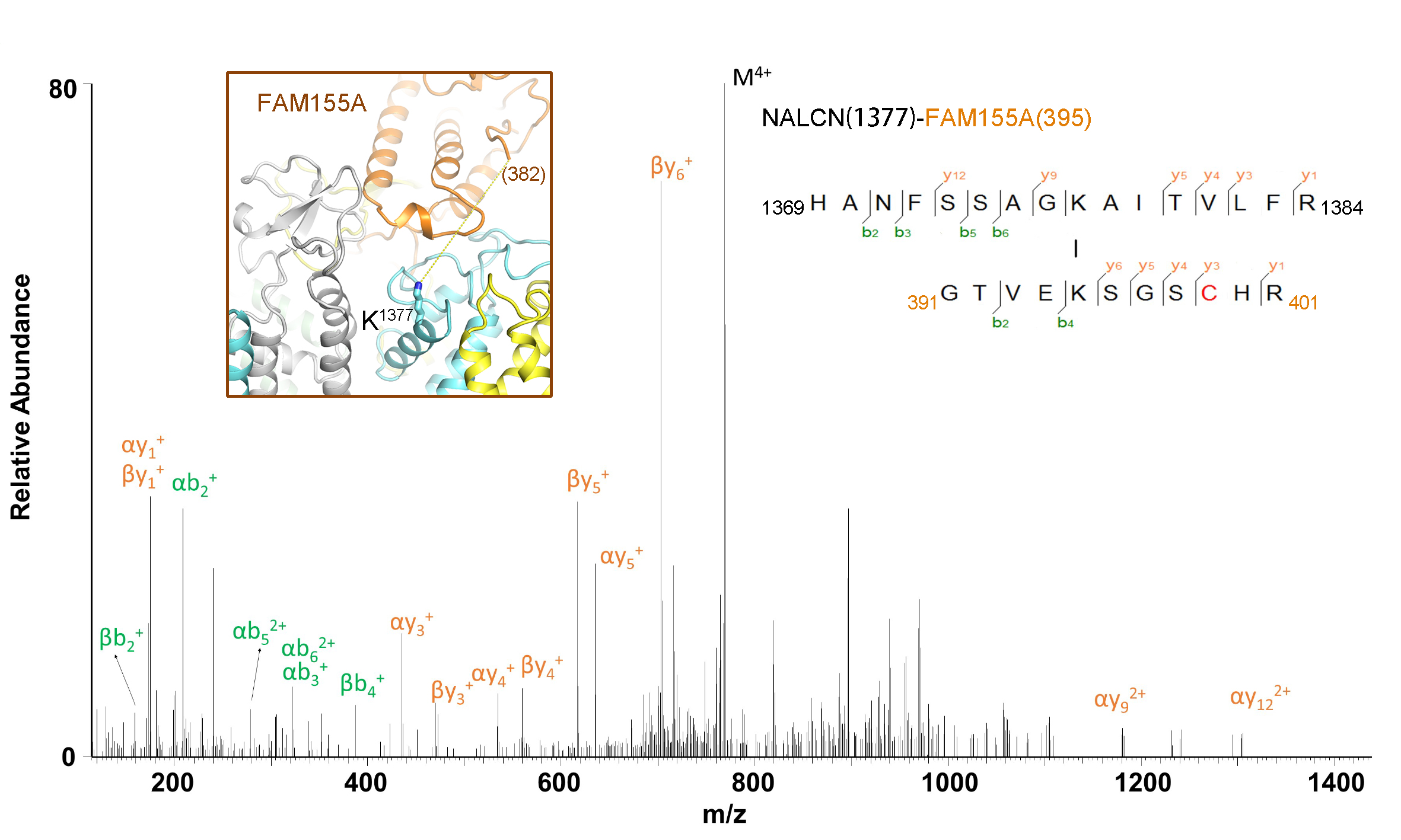

### Figure S7

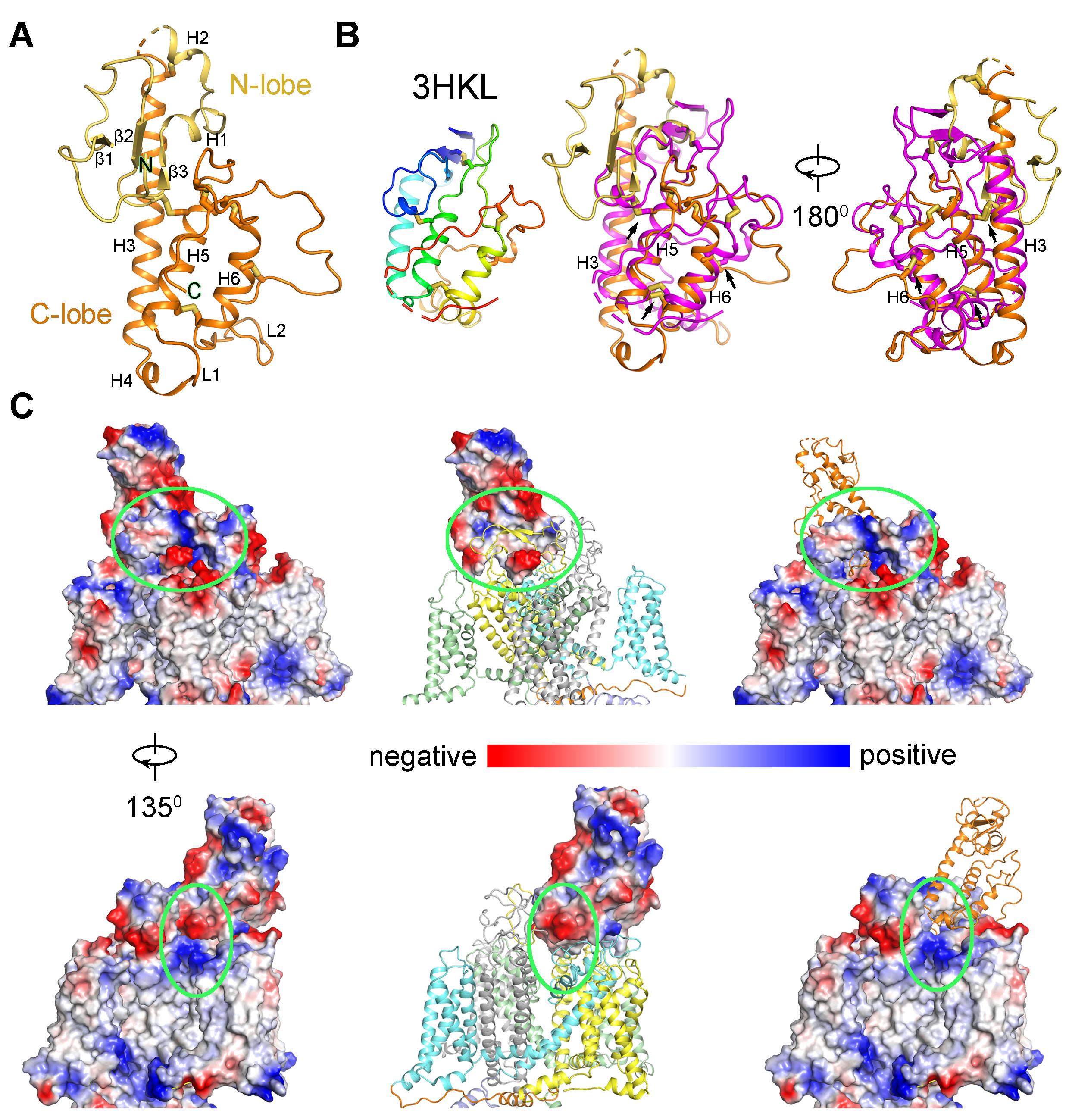

### Figure S8_p1

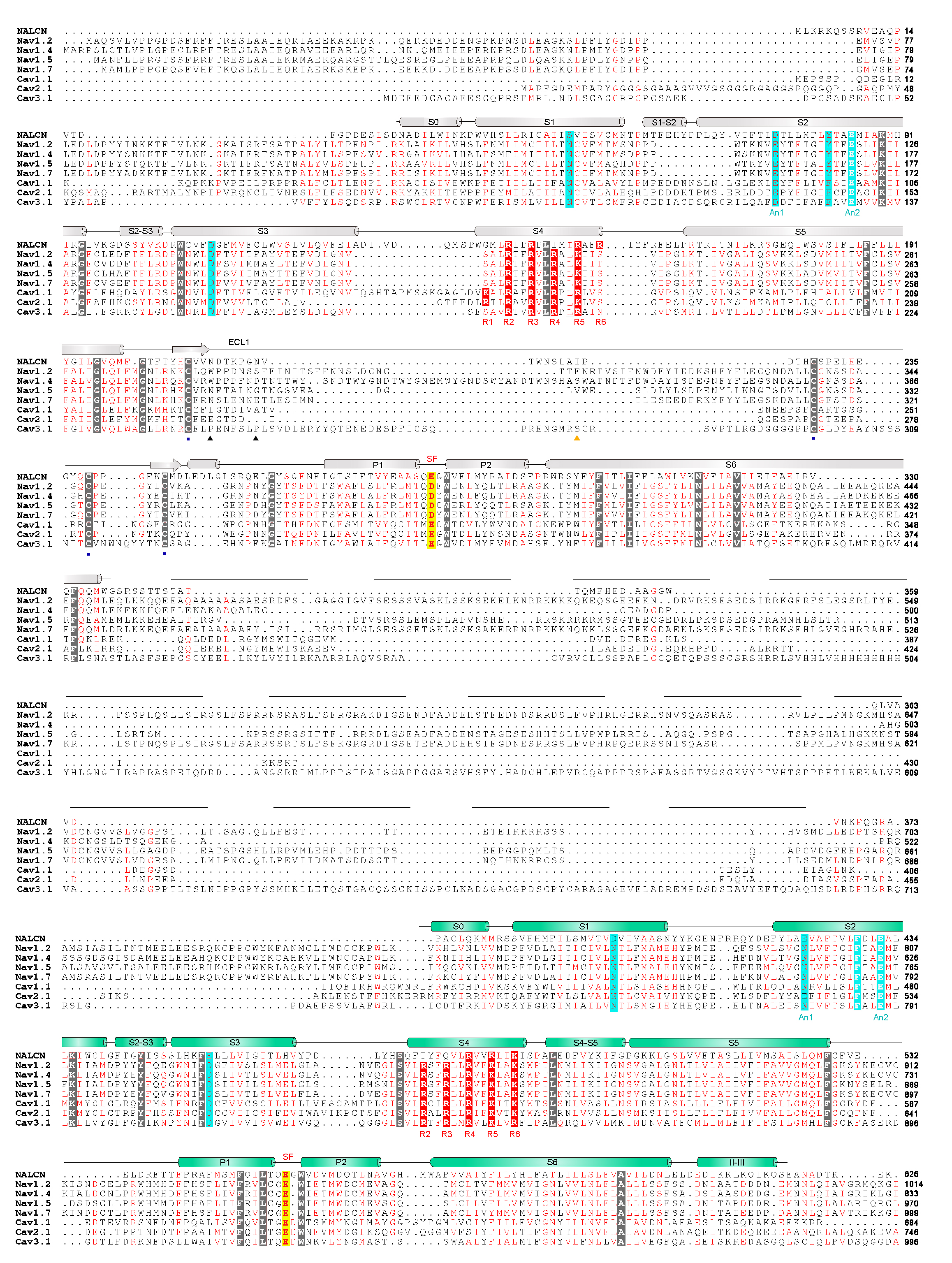

### Figure S8_p2

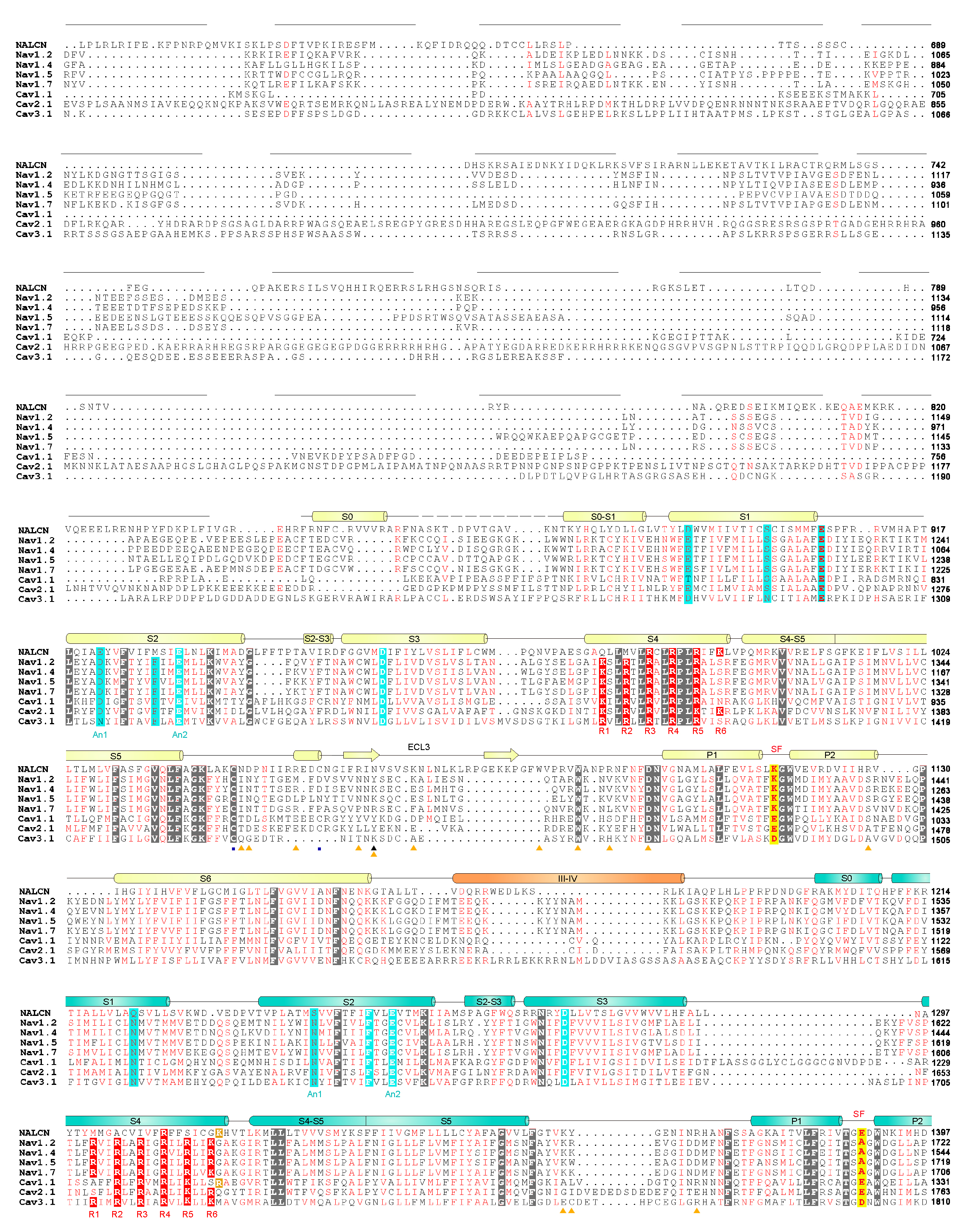

### Figure S8_p3

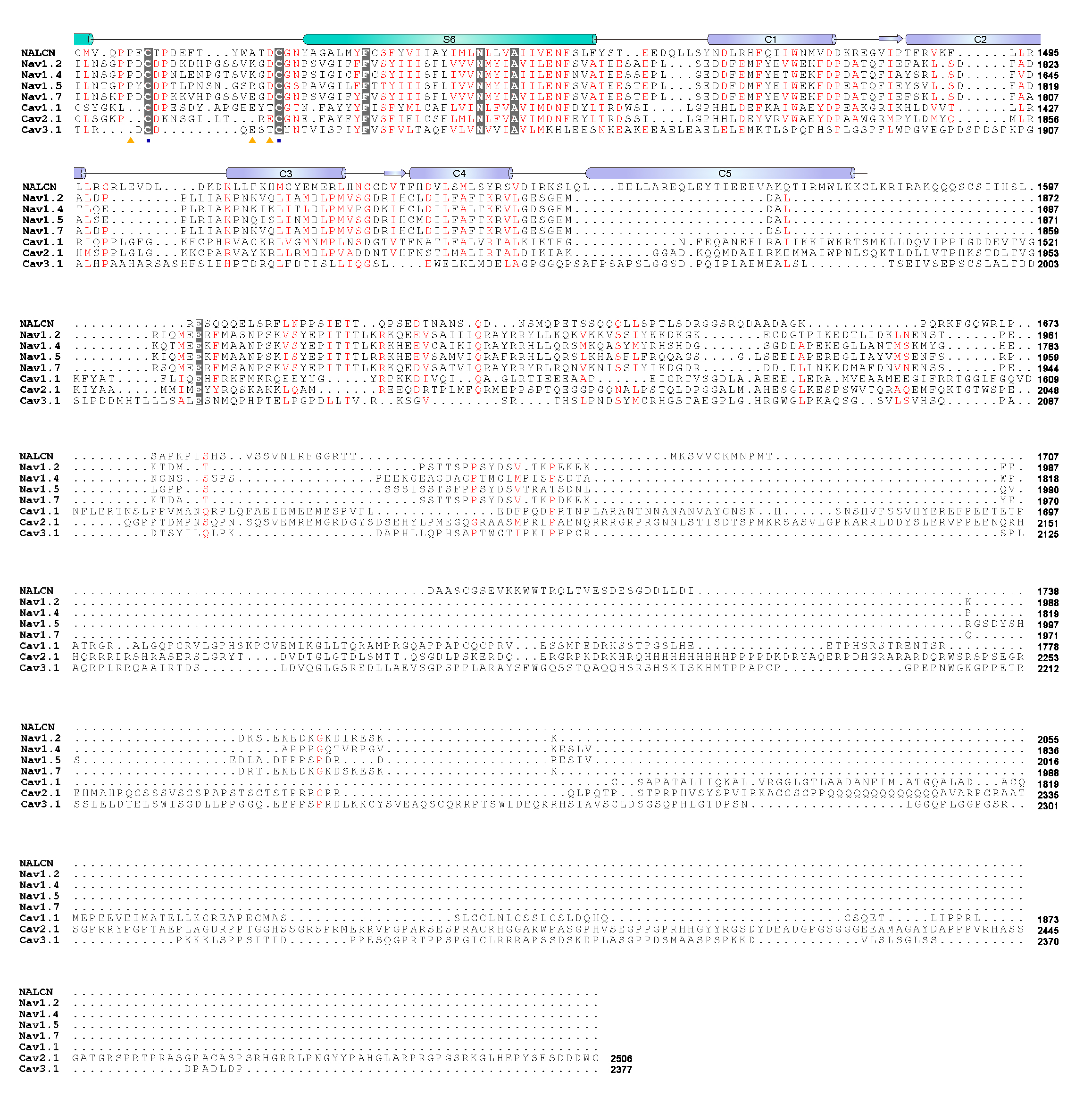

### Figure S9

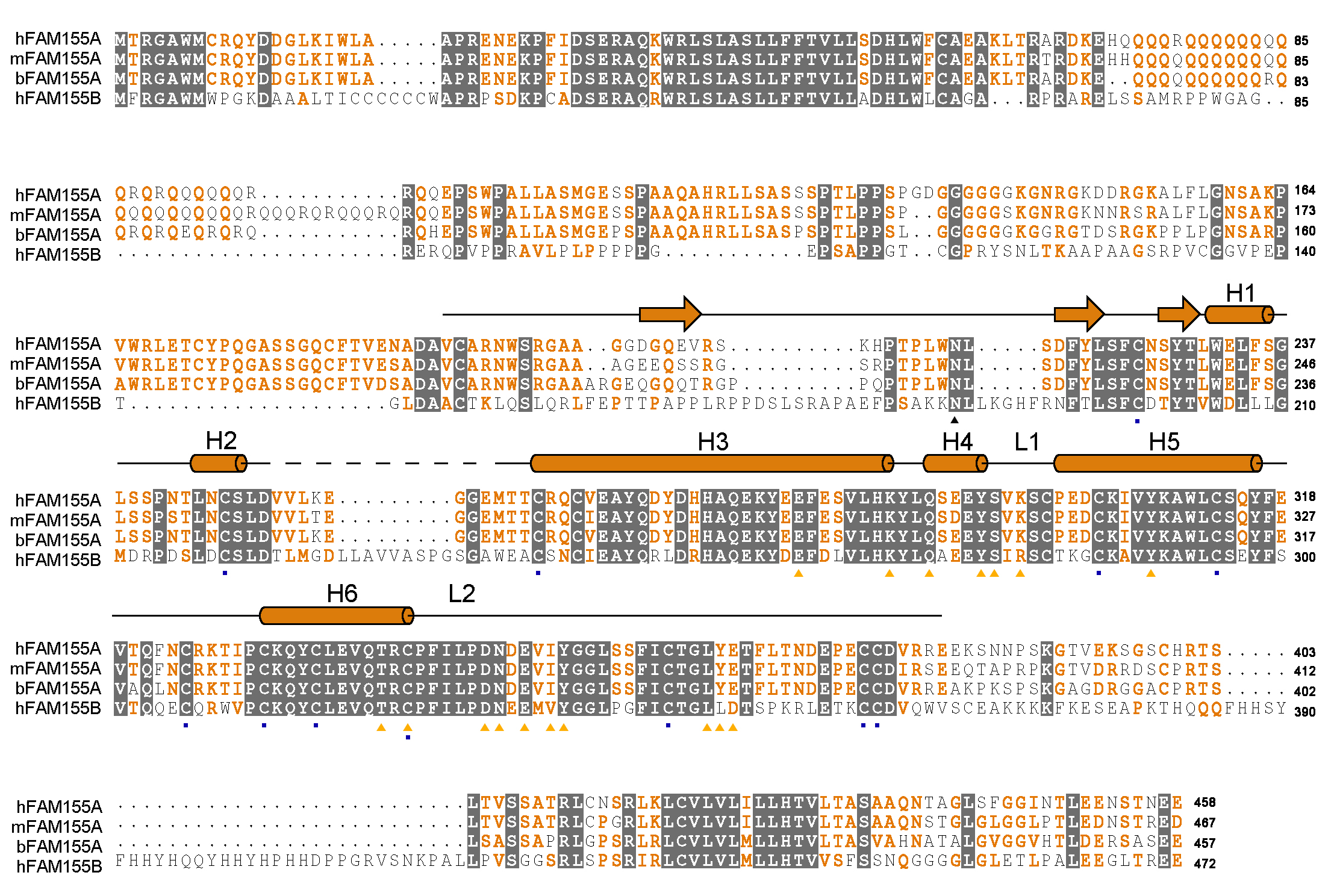

### Figure S10

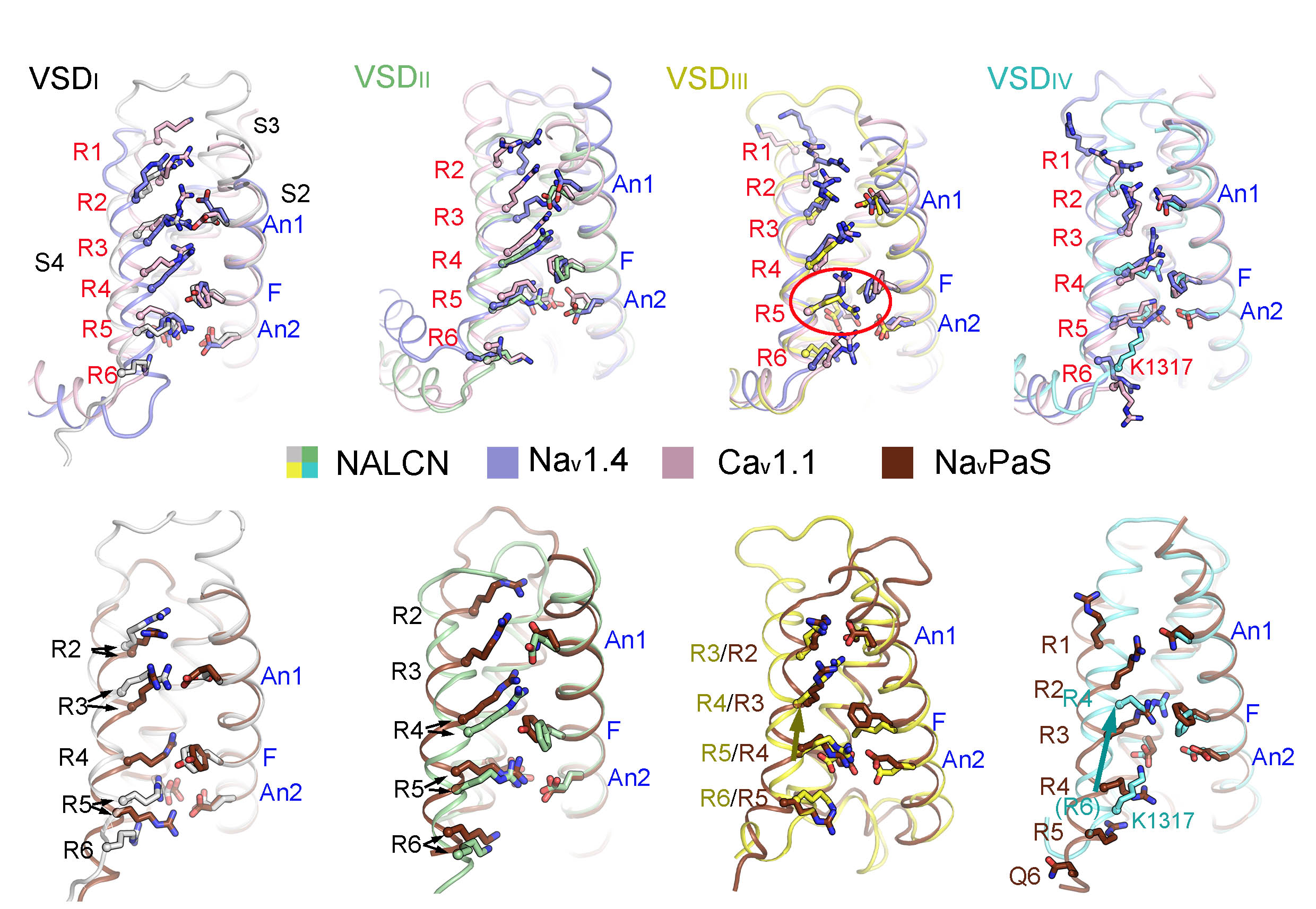

### Figure S11

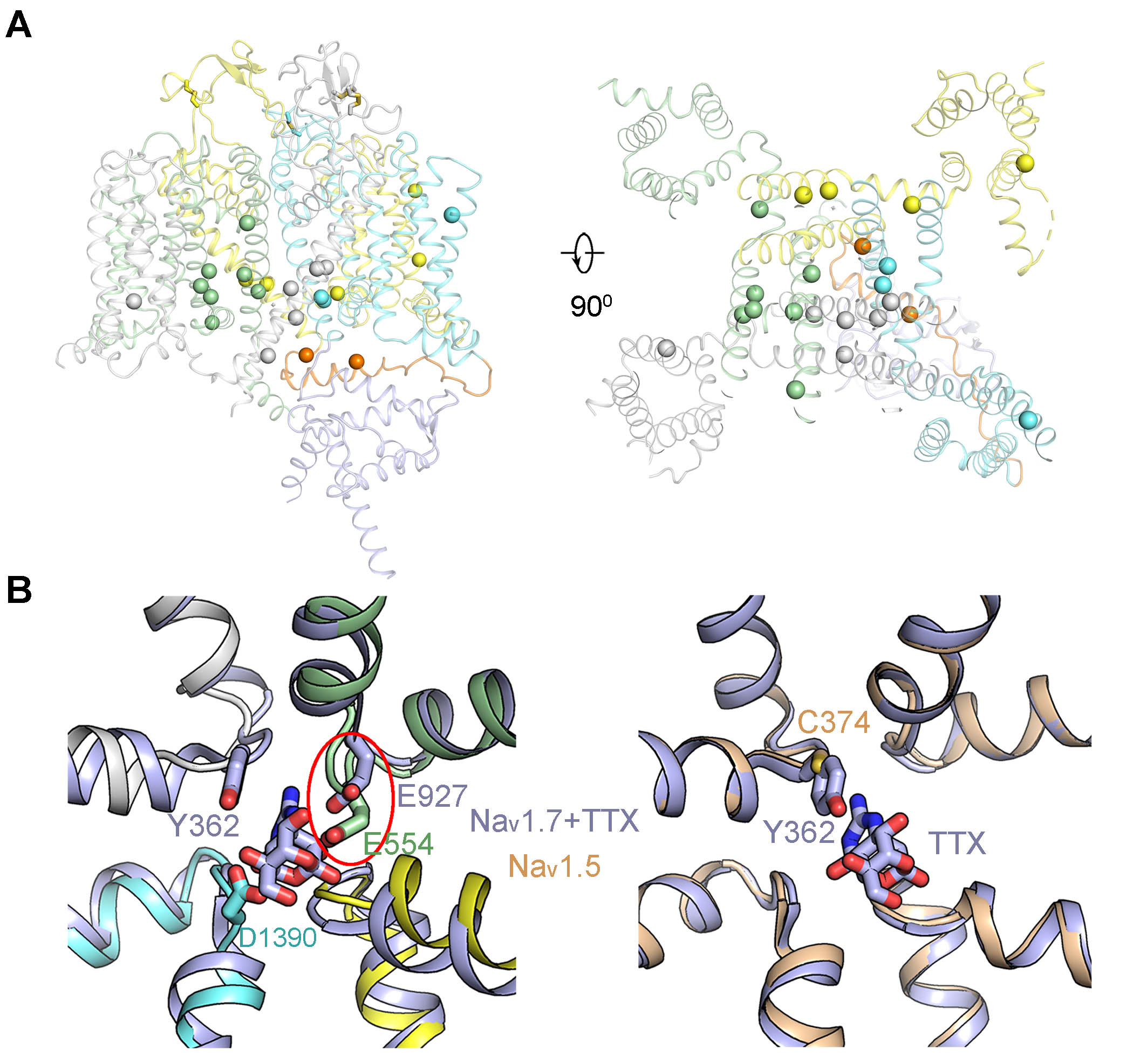
