## Supplementary material for "Structure of human sodium leak channel NALCN in complex with FAM155A": Table S1

**Table S1. Data collection and model statistics, Related to Figure 1.**

| **Data collection** |  |
| --- | --- |
| EM equipment | Titan Krios |
| Voltage (kV) | 300 |
| Detector | Gatan K3 |
| Energy filter | Gatan GIF Quantum, 20 eV slit |
| Pixel size (Å) | 1.087 |
| Electron dose (e^-^/Å^2^) | 50 |
| Defocus range (μm) | -0.5~-2.5 |
| Data set | NALCN-FAM155A complex |
| Number of images | 17,922 |
| **Reconstruction** |  |
| Software | RELION 3.0 / CryoSPARC v2 |
| Number of used Particles | 65,177 |
| Symmetry | C1 |
| Final Resolution (Å) | 3.1 |
| Map sharpening B-factor (Å^2^) | -86.5 |
| **Model building and refinement** |  |
| Model building software | Coot |
| Refinement software | Phenix |
| **Model composition** |  |
| Protein residues | 1,477 |
| Side chains | 1,463 |
| Sugar | 7 |
| Lipid | 2 |
| **Validation** |  |
| R.m.s deviations |  |
| Bonds length (Å) | 0.004 |
| Bonds Angle (˚) | 1.287 |
| Ramachandran plot statistics (%) | |
| Preferred | 89.3 |
| Allowed | 10.7 |
| Outlier | 0.0 |
