## Supplementary material for "Structure of human sodium leak channel NALCN in complex with FAM155A": Table S2

**Table S2** **Summary of contact interfaces between NALCN and FAM155A, Related to Figure 3.**

| NALCN | Residue | FAM155A | Residue |
| --- | --- | --- | --- |
| ECL_III_ | W1085 (O) | H5 | Y308 |
| ECL_III_ | W1085 | H3 | E281 |
| ECL_III_ | R1062 | H3 | E281 |
| ECL_I_ | I224 (N) | H3 | K288(O) |
| ECL_III_ | R1054 | L2 | E366 |
| ECL_III_ | W1090 (O) | L2 | L364 (N) |
| ECL_III_ | D1099 | L2 | Y365 |
| ECL_III_ | N1047 | L2 | Y353 |
| ECL_III_ | N1064 (O) | L2 | D347 (N) |
| ECL_III_ | K1069 | L2 | D347 |
| ECL_III_ | K1069 | L2 | N348 (O) |
|  | NAG (O3) | L2 | E350 |
| P2_III_ | R1127 | L2 | I352 (O) |
| ECL_III_ | R1094 (N) | L2 | G354 (O) |
| ECL_IV_ | P1403 (O) | H6 | T339 |
| ECL_III_ | D1048 | L1 | K298 |
| ECL_IV_ | A1414 (O) | L1 | K298 |
| ECL_IV_ | D1416 | L1 | S296 |
| ECL_IV_ | Y1362 (N) | L1 | Y295 (O) |
| ECL_IV_ | R1368 | L1 | Q291 |
| ECL_IV_ | K1361 | H6 | C341 (O) |

ECL: Extracellular loop
