## Supplementary material for "Structure of human sodium leak channel NALCN in complex with FAM155A": Table S3

**Table S3 Disease related mutations on human NALCN.**

| Protein | Mutations | Diseases | Structural Mapping | Reference |
| --- | --- | --- | --- | --- |
| hNALCN | R43Q | EBD | S1_I_ | (Fukai et al., 2016) |
| hNALCN | W107* | IHPRF1 | S3 _I_ | (Bramswig et al., 2018) |
| hNALCN | Q177P | CLIFAHDD | S5_I_ | (Chong et al., 2015) |
| hNALCN | W179* | IHPRF1 | *Gene shift* | (Bramswig et al., 2018) |
| hNALCN | L312I | CLIFAHDD | S6_I_ | (Chong et al., 2015) |
| hNALCN | L312V | NDH | S6_I_ | (Fukai et al., 2016) |
| hNALCN | V313G | CLIFAHDD | S6_I_ | (Chong et al., 2015) |
| hNALCN | F317C | DACH | S6_I_ | (Karakaya et al., 2016) |
| hNALCN | E327K | CLIFAHDD | S6_I_ | (Chong et al., 2015) |
| hNALCN | Y497T | ARSSHSICD | S4-S5_II_ | (Al-Sayed et al., 2013) |
| hNALCN | L509S | CLIFAHDD | S5_II_ | (Chong et al., 2015) |
| hNALCN | F512V | CLIFAHDD | S5_II_ | (Chong et al., 2015) |
| hNALCN | T513N | CLIFAHDD | S5_II_ | (Chong et al., 2015) |
| hNALCN | Y578S | CLIFAHDD | S5_II_ | (Chong et al., 2015) |
| hNALCN | L590F | CLIFAHDD | S6_II_ | (Chong et al., 2015) |
| hNALCN | V595F | DACH | S6_II_ | (Karakaya et al., 2016) |
| hNALCN | Q642* | INAD | II-III linker | (Koroglu et al., 2013) |
| hNALCN | C675L | IHPRF1 | II-III linker | (Bourque et al., 2018) |
| hNALCN | E813G | IHPRF1 | II-III linker | (Bramswig et al., 2018) |
| hNALCN | Q877N | IHPRF1 | S0-S1 _III_ | (Bramswig et al., 2018) |
| hNALCN | V891S | IHPRF1 | S1 _III_ | (Bramswig et al., 2018) |
| hNALCN | I920L | IHPRF1 | S2 _III_ | (Bramswig et al., 2018) |
| hNALCN | V956-L963del | IHPRF1 | S3 _III_ | (Bramswig et al., 2018) |
| hNALCN | V1006A | CLIFAHDD | S4-S5_III_ | (Chong et al., 2015) |
| hNALCN | R1008W | IHPRF1 | S4-S5_III_ | (Bramswig et al., 2018) |
| hNALCN | I1017T | CLIFAHDD | S5_III_ | (Chong et al., 2015) |
| hNALCN | L1019F | IHPRF1 | S5_III_ | (Bramswig et al., 2018) |
| hNALCN | V1020F | NDH | S5_III_ | (Fukai et al., 2016) |
| hNALCN | T1165P | CLIFAHDD | III-IV linker | (Chong et al., 2015) |
| hNALCN | R1181Q | NDH | III-IV linker | (Fukai et al., 2016) |
| hNALCN | Q1186* | IHPRF1 | III-IV linker | (Bramswig et al., 2018) |
| hNALCN | W1287L | IHPRF1 | S3_IV_ | (Al-Sayed et al., 2013) |
| hNALCN | R1304X | PB | S4 _IV_ | (Bourque et al., 2018) |
| hNALCN | R1384* | IHPRF1 | P1 _IV_ | (Bramswig et al., 2018) |
| hNALCN | F1427L | IHPRF1 | S6 _IV_ | (Bramswig et al., 2018) |
| hNALCN | I1445L | IHPRF1 | S6 _IV_ | (Bramswig et al., 2018) |
| hNALCN | I1446M | CLIFAHDD | S6 _IV_ | (Chong et al., 2015) |
| hNALCN | *1489ΔT* | IHPRF1 | *Gene shift* | (Al-Sayed et al., 2013) |

**Note：*** represents premature translational termination

**CLIFAHDD**: Congenital contractures of the limbs and face, hypotonia, and developmental delay;

**EBD**: Exaggerated body dends; **IHPRF1**: Hypotonia, infantile, with psychomotor retardation and characteristic facies 1; ***DACH***: Distal arthrogryposis and central hypertonia; ***NDH***: Neurodevelopmental Disease and Hypotonia; ***PB***: Periodic breathing; ***ARSSHSICD***: Autosomal recessive syndrome with severe hypotonia, speech impairment, and cognitive delay;

**INAD**: Infantile neuroaxonal dystrophy
